# Alignment-Aware Genomic Coordinate Encoding for Long-Range DNA Modeling

**DOI:** 10.64898/2026.10.10.757278

**Authors:** Xu Yang, Chenchen Qin, Fang Wang, Bing He, Yu Zhao, Jiayang Wu, Yanlin Zhang, Jianhua Yao

## Abstract

Current genomic foundation models (GFMs) largely inherit the language-modeling paradigm over contiguous token sequences, treating DNA as ordinary text. The genome, however, is a coordinate-based system of sparse signals: variants, regulatory elements, and distal gene interactions occupy a vanishing fraction of chromosome-scale sequences and are defined by their positions in the reference coordinate system rather than by linear proximity. This mismatch limits pure sequence modeling on individualized, non-contiguous genomic tasks. We present USM-AA, a genomic foundation model built on two complementary components: an alignment embedding module (AEM) that explicitly represents each token’s chromosome, strand, and comparison outcome against the reference, making variant signals directly observable; and an Alignment-Aware Rotary Position Embedding (AARoPE) that places tokens on the reference coordinate system, preserving correct positional relationships over non-contiguous inputs while supporting end-to-end training. USM-AA is further trained with a reverse-complement masked learning strategy that operates purely at the loss level, adding no parameters or architectural modification. Pretrained on large-scale genomic sequences spanning cross-species evolution and human population variation, USM-AA achieves state-of-the-art performance across multiple genomic tasks, including long-range regulatory prediction and variant-effect prediction. The source code is publicly available at https://github.com/Jimmy0743/USM-AA.

## 1 Introduction

The genome constitutes a vast and complex natural corpus composed of four nucleotides (A, T, C, and G). Unlike textual corpora in Natural Language Processing (NLP), genomic data are organized around multi-species reference genomes, where each species is associated with a shared and well-defined genomic coordinate system. The availability of large-scale, publicly accessible reference genomes across diverse species makes genomic sequences particularly well suited for training Genomic Foundation Models (GFMs) [Benegas et al., 2025b]. These models, adapted from large language model architectures, have demonstrated strong performance across a wide range of genomic tasks, such as sequence generation, chromatin modification prediction, and variant effect prediction.

Pretraining strategies for GFMs can be broadly categorized into two paradigms: masked language modeling (MLM) and autoregressive language modeling (ALM). MLM-based models, such as DNABERT2 [Zhou et al., 2024], Nucleotide Transformer (NT) [Dalla-Torre et al., 2025], and GPN-MSA [Benegas et al., 2025a], learn contextual representations by predicting randomly masked tokens within genomic sequences. In contrast, ALM-based models, including DNAGPT [Zhang et al., 2023], Evo [Nguyen et al., 2024], and GENERator [Wu et al., 2025], model sequence generation by predicting the next token in an autoregressive manner.

Most existing GFMs adopt positional encoding schemes inherited from NLP models, including AL-iBi [Press et al., 2022] in DNABERT2, rotary position embedding (RoPE) [Su et al., 2024] in Al-phaGenome [Avsec et al., 2025], and absolute positional encoding in DNABERT [Ji et al., 2021]. These schemes encode positional information defined over contiguous token sequences—relative offsets between tokens, or token indices bounded by the context length—and are effective when modeling local sequence continuity. However, none of them conveys absolute genomic coordinates, making them less suitable for non-contiguous genomic inputs such as sparsely distributed mutations or disjoint genomic regions. Concurrently, the epigenetic foundation model Pleiades [Niki et al., 2026] supplies absolute genomic coordinates through additive learned token embeddings, while leaving the index-based rotary encoding itself unchanged.

Despite their strong empirical performance, existing GFMs are fundamentally limited in their ability to represent the reference genome coordinate system. Since language models act as lossless compressors in which all task-relevant information must be captured in the learned embeddings [Delétang et al., 2023], the absence of absolute genomic coordinates during pretraining creates an unavoidable information gap. This gap has two facets: on the content side, the same *k*-mer occurs on both the reference and the alternative haplotype, so the existence of a variant is invisible at the token level; on the position side, token-order indices are a property of how the input was concatenated rather than a biological quantity, and once loci are concatenated sparsely they are uncorrelated with genomic distance while being unable to express absolute chromosome-scale position. We address these two facets with two separate mechanisms.

Sequence Alignment Map (SAM) files [Li et al., 2009] are a standard format for storing nucleotide sequence alignments against a reference genome. Each entry in a SAM file contains the sequence read, alignment position on the reference genome, CIGAR string describing matches, insertions, and deletions, as well as additional metadata such as mapping quality and strand information. This format preserves individual and reference genome variation while leveraging explicit positional information to capture nucleotide context, making it well suited for long-sequence modeling.

Inspired by the way alignment information is encoded in standard SAM files, we introduce USM-AA, a genomic foundation model that explicitly leverages reference genome coordinates and non-contiguous sequence representations. The model assigns the two quantities carried by an alignment to separate mechanisms rather than fusing them into one representation: the outcome of the comparison against the reference is encoded by the alignment embedding module (AEM) as token-level variant signal, whereas the location of the comparison is encoded by the alignment-aware rotary position embedding (AARoPE) as the rotary phase. Pretraining then uses reverse-complement masked language modeling on both contiguous and non-contiguous genomic sequences, enabling effective modeling of long-range, individual-level genomic variation.

Our contributions are organized around this decoupling of variant signal and genomic position, which yields strong performance across downstream genomic tasks:

- **Alignment Embedding Module**. Sequence-only models cannot determine whether a difference from the reference exists, because a variant base and its matching reference base are equally plausible tokens. We encode, for every token, the outcome of its comparison against the reference, covering CIGAR operation, chromosome identity, and strand, so that single-nucleotide variants (SNVs), insertions and deletions (INDELs) of both polarities, and soft clips become representable rather than merely inferable.
- **Alignment-Aware Rotary Position Embedding**. We move rotary position encoding from the sequence-index coordinate system to the reference-genome coordinate system: the rotary phase is conditioned on each token’s absolute genomic coordinate, boundary tokens are anchored to the chromosome ends, and the rotary base is enlarged to span chromosome-scale ranges. The phase difference between any two tokens equals their true genomic distance, including gaps introduced by insertions, deletions, and soft clips. Because inter-locus distances enter the attention score analytically rather than being learned from padding, the same weights apply unchanged to non-contiguous input.
- **Reverse-Complement Masked Language Modeling**: We employ a lightweight reverse-complement loss that lets the model learn the reverse-complement structure of DNA during pretraining. Unlike architecture-level reverse-complement handling in prior work [Schiff et al., 2024], RC-MLM operates purely at the objective level, adding no parameters and requiring no architectural modification, and is applicable to any encoder backbone. The model is trained to treat sequences and their reverse complements as equivalent, encouraging strand-invariant representations.

## 2 Related Work

**Alignment-Free Genomic Foundation Model** has benefited significantly from the development of Transformer architectures. One of the earliest GFMs, DNABERT [Ji et al., 2021], was derived from BERT [Devlin et al., 2019], an encoder-only Transformer model trained using MLM. Subsequent studies have largely followed the same paradigm, performing unsupervised pretraining on large-scale genomic sequences and then fine-tuning the models for downstream tasks. However, the inputs to these models are primarily limited to raw sequence information, lacking explicit alignment signals with respect to a reference genome.

**Alignment Genomic Foundation Models** incorporate sequence alignment information during training. NTv3 [Boshar et al., 2025] post-trains on functional tracks, and AlphaGenome [Avsec et al., 2025] is trained with a wide range of functional signals aligned onto the reference genome. In both models, however, this alignment only connects experimental measurements to DNA sequences during training, and is not used as input at inference time. In contrast, USM-AA treats the alignment against the reference genome as a first-class input throughout pretraining and inference, encoding each token’s comparison outcome through the alignment embedding module and its reference coordinate through AARoPE.

**Positional Embedding** is a core component of Transformer architectures, enabling models to encode token order. Common approaches include absolute position embeddings, sinusoidal encodings [Vaswani et al., 2017], ALiBi [Press et al., 2022], and RoPE [Su et al., 2024]. Absolute and sinusoidal embeddings represent positions explicitly, while ALiBi and RoPE encode relative positional information. These methods have been widely adopted in NLP and large-scale genomic models such as DNABERT2 [Zhou et al., 2024] and AlphaGenome [Avsec et al., 2025]. However, they are primarily designed for continuous sequences and do not account for alignment-aware or reference-based positional structures.

## 3 Methodology

### 3.1 Overview

As illustrated in Fig. 1, USM-AA is a Transformer-based encoder model pretrained on a combination of the human reference genome, population-scale genomes from the 1000G [Abecasis et al., 2010], and reference genomes from multiple species. Input sequences are first aligned to hg38 to obtain alignment attributes (Sec. 3.2), which are encoded by an alignment-aware representation module (Sec. 3.3) and processed by Transformer encoder layers (Sec. 3.4). The model is pretrained with RC-MLM (Sec. 3.5). As a result, USM-AA captures both sequence content and coordinate-aware genomic structure, and is suited to interpret genomic sequences in the context of the reference genome.

**Figure 1:**
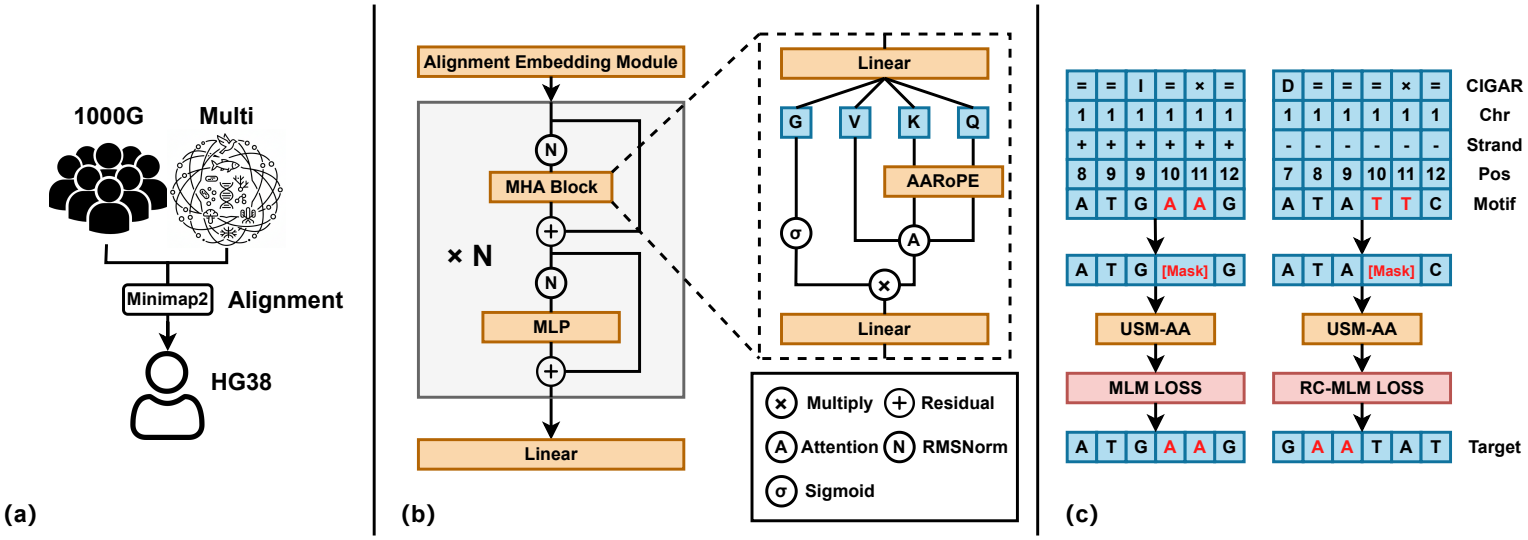
Overview of the USM-AA architecture. (a) Pretraining data sources include the human reference genome hg38, population-scale genomes from the 1000 Genomes Project, and reference genomes from multiple species. (b) USM-AA builds an alignment-aware representation comprising an alignment embedding module and AARoPE, followed by *N* Transformer layers with gated attention, enabling the processing of both contiguous and non-contiguous sequences. (c) The model is trained with a strand-aware masked language modeling objective. For sequences aligned to the forward strand, standard MLM is applied, whereas for sequences aligned to the reverse strand, the model is required to predict the reverse-complement nucleotides at the masked positions.

### 3.2 Data Alignment and Tokenization

Instead of relying solely on sequence information, we align all pretraining and fine-tuning datasets to the human reference genome (hg38) [Schneider et al., 2017] using Minimap2 [Li, 2018]. This alignment provides, for each token, its absolute genomic coordinate on the corresponding chromosome, strand information, and the CIGAR value, capturing variations such as SNVs, INDELs, and soft clips relative to the reference sequence. Alignable sequences are processed using alignment-aware training, whereas unalignable sequences are incorporated through sequence-only training.

For tokenization, we adopt a dynamic k-mer strategy: sequences are primarily segmented into non-overlapping 6-mers, while residual nucleotides that cannot form a complete 6-mer are tokenized using shorter k-mers. Further details are provided in Appendix B.6.

### 3.3 Alignment-Aware Representation

#### 3.3.1 Alignment Embedding Module

Each token carries an alignment tuple *a*_*i*_ = (*ℓ*_*i*_, *c*_*i*_, *s*_*i*_, *g*_*i*_): its genomic coordinate *ℓ*_*i*_, chromosome *c*_*i*_, strand *s*_*i*_, and CIGAR operation *g*_*i*_. The alignment embedding module fuses these attributes additively:

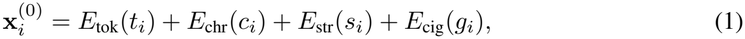

where *E*_chr_, *E*_str_, *E*_cig_ are learnable chromosome, strand, and CIGAR embeddings (see Appendix B.5 for the encoding scheme). The coordinate *ℓ*_*i*_ carries no embedding here; it enters the model exclusively through AARoPE (Sec. 3.3).

#### 3.3.2 AARoPE

AARoPE optimizes RoPE [Su et al., 2024] by conditioning the rotary phase on the absolute reference-genome coordinate *ℓ*_*i*_ instead of the token index.

For token *i* with genomic coordinate *ℓ*_*i*_ on chromosome *c*_*i*_ in the reference coordinate system, the phases and rotated queries/keys are

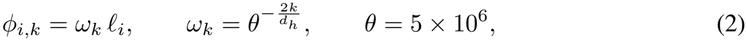

and the queries and keys are rotated by the standard block-diagonal rotary matrix assembled from these phases [Su et al., 2024], i.e., 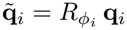 and 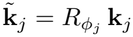, where **q**_*i*_ = **x**_*i*_*W*_*Q*_, **k**_*j*_ = **x**_*j*_*W*_*K*_ with **x**_*i*_ denoting the token representation at the current layer.

##### Absolute position

Boundary tokens (*ℓ*_BOS_/*ℓ*_EOS_, where BOS and EOS denote the begin- and end-of-sequence tokens) are anchored to the chromosome ends (*L*_*c*_: chromosome length):

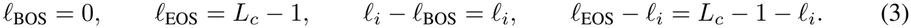

Both distances to the chromosome ends are therefore recoverable from the rotary phase, so the encoding is absolute at chromosome scale.

##### Relative position

The phase difference depends only on the coordinate difference:

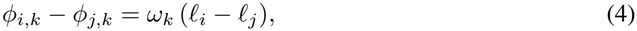

which enters the attention score as

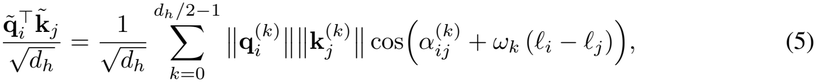

where 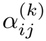 is the initial angular offset in plane *k*. The attention score thus encodes the true genomic distance; meanwhile, *c*_*i*_ is injected via the chromosome embedding into the rotated Q/K (Eq. 1), distinguishing identical coordinates on different chromosomes.

### 3.4 Transformer Encoder

We train two scales of USM-AA models: USM-AA-100M (approximately 122M parameters) and USM-AA-1B (approximately 1.16B parameters). As shown in Fig. 1(b), the encoder consists of *N* stacked Pre-Norm Transformer layers [Zhang and Sennrich, 2019] with Gated Attention [Qiu et al., 2025] for fine-grained feature selection; the complete architectural details are provided in Appendix B.1.

### 3.5 Reverse-Complement Masked Language Modeling

DNA exists as a double helix with two complementary strands running in opposite directions. To exploit this biological structure, we employ a lightweight reverse-complement loss that lets the model learn the correspondence between forward and reverse complement strands.

Notably, our approach operates purely at the *objective* level and is orthogonal to architecture-level reverse-complement handling: Caduceus [Schiff et al., 2024] enforces RC equivariance by weight-sharing across the forward and reverse strands within the network architecture, whereas RC-MLM is a lightweight training loss that adds no parameters and is applicable to any encoder backbone without architectural modification.

Specifically, given a token sequence **t** = [*t*_1_, *t*_2_, …, *t*_*n*_] aligned to strand *s* with mask positions *ℳ*, the model is trained to predict the strand-consistent target: for forward-strand inputs (*s* = 0), the standard MLM target; for reverse-strand inputs (*s* = 1), the reverse complement of the original sequence:

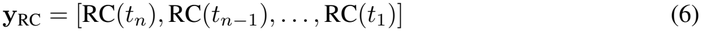

where RC(·) maps each nucleotide to its complement (A↔T, C↔G).

The combined pretraining objective is:

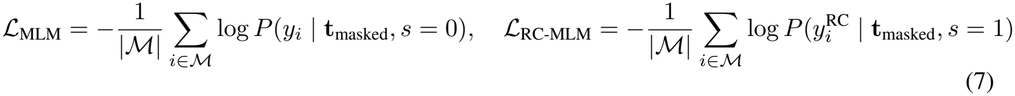

where *s* ∈ {0, 1} denotes the strand indicator, and *i* ∈ *ℳ* denotes the index of a masked token.

This hybrid loss enables the model to learn strand-aware representations, capturing the inherent symmetry of the DNA double helix and facilitating downstream tasks that involve both strands of the genome.

### 3.6 Data

During pretraining, we adopt a mixed-data strategy that integrates three complementary genomic data sources to support training on both contiguous and non-contiguous genomic sequences: (1) the human reference genome (hg38, ~ 3.2 Gbp), which serves as the canonical coordinate system for alignment-aware pretraining; (2) the DNABERT2 multi-species corpus (~32.5 Gbp from 135 species), which helps minimize variability arising from differences in pretraining data; and (3) the 1000 Genomes Project [Abecasis et al., 2010], from which we construct variant-based sequences using SNVs and INDELs from 3,202 individuals (~31.7 Gbp of alternative-allele content).

### 3.7 Training Details

Both model scales are pretrained with AdamW under a cosine learning rate schedule, using BFloat16 mixed precision and DeepSpeed-based distributed training. The complete set of pretraining hyperparameters is provided in Appendix B.4.

## 4 Experiments

### 4.1 Downstream tasks

We evaluate USM-AA at two scales (100M and 1B parameters), and primarily report results for the 1B model. Downstream tasks are trained with average pooling and a linear classification head under full-parameter fine-tuning. Our evaluation focuses on two problems targeted by USM-AA, long-range regulatory modeling and variant signal processing, followed by standard genomic model benchmarks that assess general-purpose sequence understanding.

#### 4.1.1 Expression Quantitative Trait Loci Prediction

Expression quantitative trait loci (eQTL) are nucleotide variants that influence the expression of one or more genes [Nicolae et al., 2010], and predicting their effects from sequence alone requires modeling gene regulation across long genomic distances. We adopt the long-range eQTL task from DNALONGBENCH [Cheng et al., 2025], whose dataset covers nine human cell types and consists of reference–alternative sequence pairs that differ only at the variant site; on this task, the performance of existing DNA foundation models remains unsatisfactory.

In the original protocol, each sample is padded with N bases to a fixed length of 450,000 bp, so prior models must consume an ultra-long contiguous sequence in which the discriminative signal occupies only two isolated loci; such a single-base variant signal is likely to be diluted during long-sequence modeling. USM-AA instead consumes only two short windows centered on the variant and the transcription start site (TSS), with no padding or masking, and relies on AARoPE to encode their absolute genomic coordinates. eQTL is thus the setting where the two components of the alignment-aware representation act jointly: the alignment embedding module represents the variant as a token that mismatches the reference genome, while AARoPE encodes the variant–TSS separation, so the reference and alternative conditions are compared within one coordinate-aware representation and adapted end-to-end.

We compare USM-AA against the baseline models of the DNALONGBENCH benchmark, fine-tuning on the dataset splits provided in the original benchmark. As shown in Fig. 2 and Table 8, USM-AA-1B achieves an average area under the receiver operating characteristic curve (AUROC) of 0.897 across the nine cell types, substantially surpassing the best baseline model (the Expert Model, 0.681). This result highlights the modeling advantages of AARoPE and AEM on tasks involving long-range variant–gene regulatory relationships: AEM renders the single-base variant explicitly observable at the token level, while AARoPE preserves the exact genomic separation between the variant and the TSS, enabling the two conditions to be compared within one coordinate-aware representation. By decoupling variant signal from genomic position, USM-AA offers a promising foundation for scaling alignment-aware modeling to individual-level regulatory genomics. Baseline configurations, dataset composition, exact window design, and full results are provided in Appendix C.

**Figure 2:**
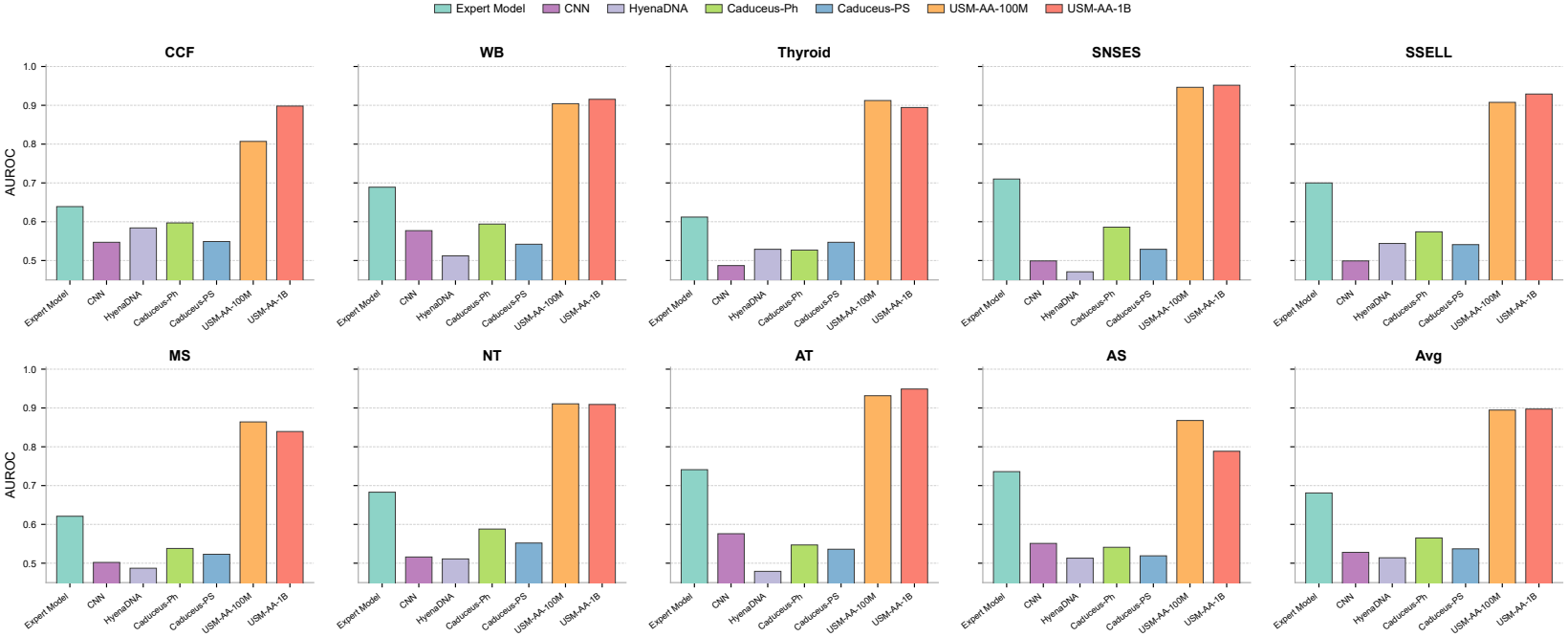
Per-cell-type eQTL prediction performance. AUROC of each model across the nine human cell types and their average, ordered as in Table 8. The USM-AA variants rank first or second in all nine cell types.

#### 4.1.2 Enhancer Target Gene Prediction

Enhancer-target gene prediction (ETGP) predicts whether a given enhancer regulates a target gene [Fulco et al., 2019], with interaction distances up to 450 kb, and therefore evaluates whether the model can capture long-range regulatory interactions over non-contiguous inputs: unlike eQTL, no comparison against a variant is involved, and AARoPE alone must carry the interaction signal across the two segments. We adopt the ETGP task from DNALONGBENCH [Cheng et al., 2025], whose original protocol likewise pads each sample to 450,000 bp; USM-AA instead consumes only the enhancer region and a short window centered on the TSS. We compare against the DNALONG-BENCH baseline models, including the ABC-based Expert Model that additionally incorporates DNase-seq, H3K27ac ChIP-seq, and Hi-C data. As shown in Table 1, USM-AA outperforms all baselines. Dataset details are provided in Appendix D.

**Table 1:** Enhancer-target gene prediction performance on DNALONGBENCH [Cheng et al., 2025]. Best scores are bold, second best are underlined.

| Model | USM-AA-1B | USM-AA-100M | Expert Model | HyenaDNA | Caduceus-PH | Caduceus-PS | CNN |
|---|---|---|---|---|---|---|---|
| AUROC | **0.952** | <u>0.934</u> | 0.926 | 0.828 | 0.826 | 0.821 | 0.797 |

#### 4.1.3 TCGA Cancer Classification

Beyond regulatory signals, we leverage the non-contiguous modeling capability of USM-AA to represent an individual genome at the variant level and construct an individual-level cancer-type classification task. Using the TCGA Pan-Cancer Atlas [Hoadley et al., 2018], which profiles 33 cancer types, we represent each individual as the set of their germline-filtered somatic variants, including SNVs and INDELs, sparsely distributed across all chromosomes. This yields a 33-way classification task in which the input is not a contiguous sequence but a sparse, genome-wide collection of variant signals. Such a format is natural for USM-AA, which takes all variants of an individual as a single non-contiguous input: AEM attaches the alignment attributes of each variant, including its chromosome, strand, and comparison outcome, while AARoPE encodes its absolute genomic coordinate, enabling joint modeling of genome-wide variant signals. In contrast, prior genomic foundation models require contiguous sequence inputs and cannot directly represent individual genomes in this form. Given the severe class imbalance across the 33 cancer types, we adopt macro-averaged F1 as the primary evaluation metric, as it weights all classes equally and better reflects performance on rare cancer types than accuracy.

As shown in Table 2, USM-AA achieves the best performance among all methods, demonstrating its capability to process genome-wide sparse variant signals through alignment-aware modeling. Implementation details are provided in Appendix E.

**Table 2:** TCGA Pan-Cancer classification performance across 33 cancer types. Best scores are bold, second best are underlined.

| Model | F1 | Accuracy | Precision |
|---|---|---|---|
| USM-AA-1B | **0.379** | **0.436** | **0.398** |
| USM-AA-100M | <u>0.343</u> | <u>0.413</u> | <u>0.360</u> |
| Transformer | 0.319 | 0.399 | 0.330 |
| XGBoost | 0.246 | 0.345 | 0.289 |
| CNN | 0.208 | 0.298 | 0.236 |
| Logistic Regression | 0.113 | 0.234 | 0.269 |

#### 4.1.4 NT benchmarks

To evaluate the performance of USM-AA on general genomic modeling tasks, we benchmark it on 18 downstream tasks from the NT benchmark [Dalla-Torre et al., 2025], which encompass a diverse set of human genomic prediction problems, including histone modification prediction, cisregulatory element annotation, and splice site recognition. Following prior work, we adopt MCC as the primary evaluation metric, and all models are evaluated using 10-fold cross-validation. Baseline comparisons include DNABERT2, NT, NTv3, Enformer [Avsec et al., 2021], HyenaDNA [Nguyen et al., 2023], and SPACE [Yang et al., 2025]. Detailed evaluation protocols and results are provided in Appendix F.3.

As summarized in Table 3, USM-AA achieves top performance in two of the three main task categories and attains the best average MCC across all 18 tasks. In the detailed results provided in Appendix Table 13, USM-AA attains the highest MCC on 11 out of 18 tasks, outperforming all baseline models. Performance gains are especially pronounced for tasks involving Chromatin Profiles and Regulatory Elements prediction.

**Table 3:** Average Matthews correlation coefficient (MCC) performance on the NT benchmark by task category. Avg denotes the average MCC over all 18 tasks. Best results are shown in **bold**, and second-best results are underlined.

| Model | #Params | Chromatin | Regulatory | Splicing | Average |
|---|---|---|---|---|---|
| DNABERT2 | 117M | 55.7 | 66.0 | 85.1 | 63.5 |
| HyenaDNA-32KB | 3.9M | 50.3 | 60.3 | 87.7 | 59.3 |
| NT-HumanRef | 500M | 53.1 | 65.9 | 94.4 | 63.6 |
| NT-Multispecies | 2.5B | 54.9 | 69.8 | **96.4** | 65.9 |
| NTv3-pre | 650M | 55.0 | 71.1 | 93.6 | 65.9 |
| NTv3-post | 650M | 60.9 | 74.4 | 85.4 | 68.7 |
| Enformer | 252M | 55.8 | 69.8 | 75.6 | 63.0 |
| SPACE | 588M | 59.0 | 71.8 | 91.7 | 68.0 |
| USM-AA | 122M | 62.2 | 74.5 | 94.2 | 70.9 |
| USM-AA | 1.16B | **65.6** | **75.1** | 95.1 | **73.1** |

### 4.1.5 GUE and Genomic Benchmarks

We further evaluate USM-AA on two additional benchmarks for broader genomic understanding and cross-species generalization: the Genomic Understanding Evaluation (GUE) benchmark [Zhou et al., 2024], which covers seven task categories spanning human, mouse, yeast, and virus, and the Genomic Benchmarks (GB) [Grešová et al., 2023], which consists of nine regulatory-sequence classification datasets across four organisms. Both evaluations follow the same training and evaluation protocol as prior work, with baseline results taken from the original studies for fair comparison. On GUE, as shown in Table 16, USM-AA attains the highest performance on four of the seven major task categories, and its superior performance on the EMP tasks in yeast demonstrates that it maintains excellent performance even on sequences from other species. On GB, as shown in Table 19, USM-AA achieves the best performance on six of the nine tasks, further confirming its generalization capability across diverse organisms and regulatory settings. Detailed training procedures and complete results are provided in Appendices G and H.

### 4.2 Ablation study

#### 4.2.1 Component Analysis

To analyze the contribution of each component, we conduct an ablation study on the NT benchmark. As shown in Table 4, we systematically remove AARoPE, alignment embeddings (AEM), and RC-MLM in different combinations. “w/o AARoPE” uses relative positional encoding instead of AARoPE while retaining alignment embeddings. “w/o AEM” removes all alignment embedding information. “w/o RC” removes RC-MLM during pretraining.

**Table 4:**
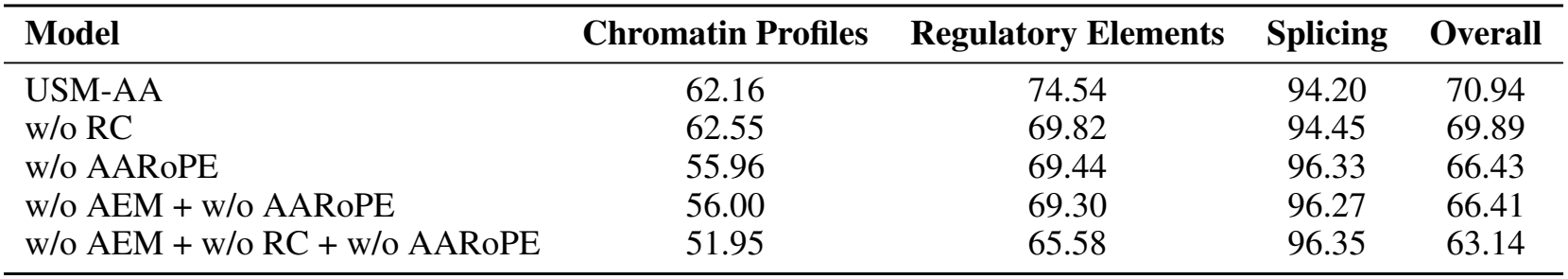
Ablation study on USM-AA components.

The results show that removing any single component leads to a noticeable drop in overall performance, confirming that each module contributes to the overall effectiveness. Furthermore, removing multiple components simultaneously causes progressively larger degradation, indicating strong synergy among the proposed modules.

#### 4.2.2 Position Embedding

To evaluate the effectiveness of different positional encoding methods for aligned genomic sequences, we compare model perplexity on reference sequences using AARoPE, sinusoidal embeddings, ALiBi, and linear embeddings.

Models are pretrained on a corpus with 50% contiguous and 50% randomly sampled non-contiguous sequences from the reference genome, and positional encoding methods are evaluated via perplexity on the evaluation set. As shown in Fig. 3(a), the parameter *K* denotes the downsampling ratio, where each sequence is constructed by uniformly sampling 1*/K* of the tokens from the original sequence and concatenating them. As *K* increases, corresponding to increasingly sparse non-contiguous sequences, AARoPE consistently outperforms the other positional encoding methods, demonstrating its superior robustness to non-contiguous sequences. Additional details on positional embeddings and perplexity computation are provided in Appendix A.1 and Appendix I.2. Robustness of USM-AA to non-contiguous downstream inputs is further evaluated in Appendix I.4.

**Figure 3:**
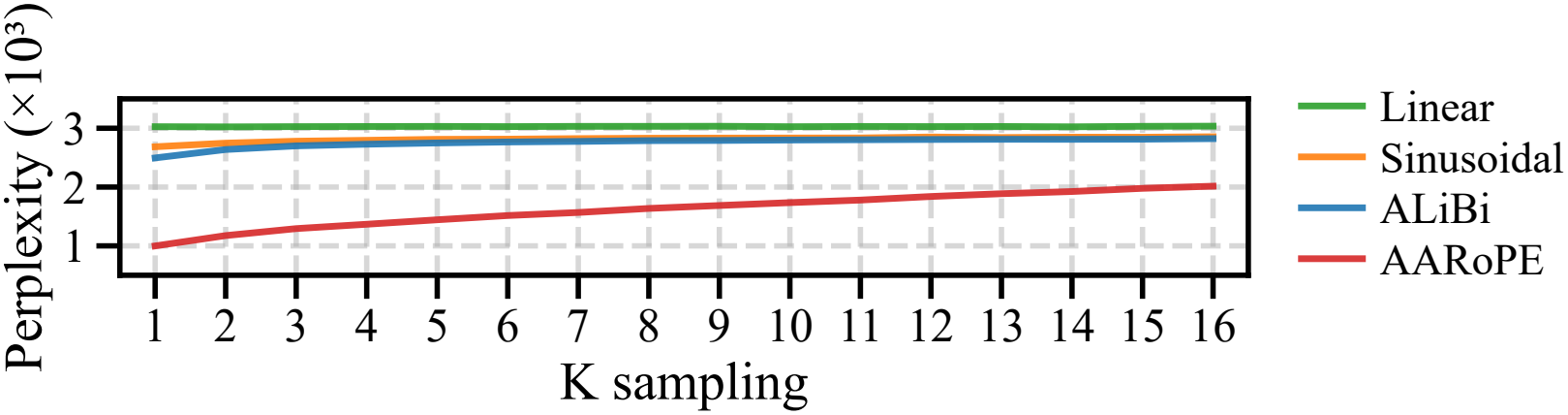
Perplexity of different positional encoding methods evaluated on non-contiguous sequences with varying sparseness. The parameter *K* denotes the downsampling ratio, where each sequence is constructed by uniformly sampling 1*/K* of the tokens from the original sequence and concatenating them.

## 5 Conclusion

We introduce USM-AA, a genomic foundation model pretrained on aligned genome data using a unified alignment-aware representation that couples AARoPE with alignment embeddings. USM-AA effectively models both contiguous and non-contiguous genomic inputs and achieves state-of-the-art performance across a range of downstream tasks. Our findings underscore the importance of alignment-aware representations in genomic sequence modeling and suggest a promising direction for future genomic foundation models.

## Limitations

While USM-AA performs well on human genomic tasks, it is pretrained using only the human reference genome as the coordinate system. For multi-species sequences that cannot be aligned to this reference, alignment attributes are unavailable and the model falls back to sequence-only training during pretraining. Additionally, the current framework relies on a single linear reference genome, which cannot fully capture the structural diversity present in human populations. The emerging pangenome paradigm, which represents genomic variation as a graph rather than a linear sequence, offers a more comprehensive model of population-level diversity. Future work could extend USM-AA to multi-species pretraining, incorporate multiple reference coordinate systems, and explore graph-based pangenome representations to improve cross-species applicability and better capture structural variants.

### Ethical Considerations

This work uses only publicly available, de-identified datasets, including the human reference genome, the 1000 Genomes Project, GTEx, TCGA Pan-Cancer Atlas, and standard genomic benchmarks. No new human subjects data were collected, and no individual-level re-identification is performed or enabled by our methods.

## A Theoretical Formulation

### A.1 Evaluation Formulation

#### A.1.1 Perplexity

Perplexity (PPL) is a commonly used metric for evaluating the performance of generative language models, such as GPT [Radford et al., 2018], on a given corpus. Since our models are trained using a RC-MLM objective, we adopt the PPL computation method introduced in NT [Dalla-Torre et al., 2025], which is adapted for masked token reconstruction.

Specifically, given a sequence *t* and a set of masked positions *ℳ*, the model predicts the probabilities of the true tokens at the masked positions. The average cross-entropy loss over the masked positions is computed as

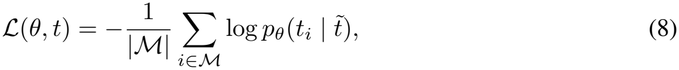

where 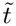 is the masked sequence and 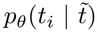 denotes the probability assigned by the model to the true token *t*_*i*_ at position *i*. The PPL is then defined as the exponential of this loss:

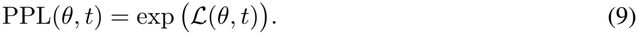

Perplexity thus measures how well the model can reconstruct masked tokens. Compared to accuracy, which only considers whether the most probable token matches the true token, PPL also accounts for the predicted probability distribution, making it a more refined measure of reconstruction performance.

#### A.1.2 Matthews Correlation Coefficient

MCC is a widely used evaluation metric for assessing the quality of binary and multiclass classification models in machine learning. MCC can be interpreted as a correlation coefficient between the predicted labels and the ground-truth labels, with values ranging from −1 to +1. A value of +1 indicates a perfect prediction, 0 corresponds to random prediction, and −1 represents a completely inverse prediction.

For binary classification, MCC is defined as

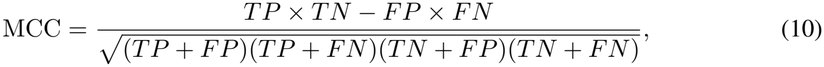

where *TP, TN, FP*, and *FN* denote the numbers of true positives, true negatives, false positives, and false negatives, respectively.

For multiclass classification with *N*_*c*_ classes and *N* samples, MCC can be computed from the confusion matrix 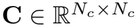 as

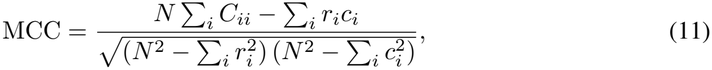

where *r*_*i*_ = Σ_*j*_ *C*_*ij*_ and *c*_*i*_ = Σ_*j*_ *C*_*ji*_ denote the total number of true and predicted samples for class *i*, respectively.

#### A.1.3 Accuracy (ACC)

Accuracy measures the proportion of correctly predicted samples:

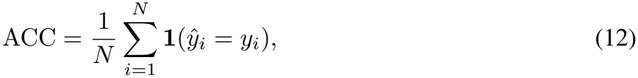

where *ŷ*_*i*_ and *y*_*i*_ are the predicted and true labels for sample *i*, and **1**(·) is the indicator function.

#### A.1.4 F1-score

The F1-score is the harmonic mean of precision and recall:

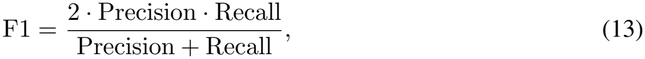

with

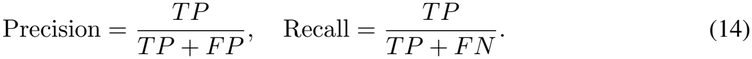

For multiclass classification, we adopt the macro-average F1, which computes the F1-score for each class independently and then averages over all classes.

## B Pretraining

### B.1 Model Architecture

USM-AA-100M consists of 12 layers with 12 attention heads and a hidden dimension of 768, resulting in approximately 122M parameters, while USM-AA-1B scales up to 36 layers with 32 attention heads and a hidden dimension of 1536, with approximately 1.16B parameters. Each encoder layer employs a Pre-Norm architecture with RMSNorm [Zhang and Sennrich, 2019] for normalization. The self-attention module uses the Gated Attention mechanism described below. The attention computation is accelerated using Flash Attention [Dao et al., 2022]. The feed-forward network uses the SwiGLU [Shazeer, 2020] activation function with a hidden dimension of 4× the embedding dimension. The pretrained weights of both models are publicly available on Hugging Face at https://huggingface.co/JimmyYang2025/usmaa_100m and https://huggingface.co/JimmyYang2025/usmaa_1b.

### B.2 Gated Attention

We employ Gated Attention [Qiu et al., 2025] to enable the model to learn more fine-grained feature selection. Specifically, the input is projected through a linear layer into four components: *Q, K, V, G*, where *G* serves as the gating vector. The attention output is element-wise multiplied with the sigmoid activation of the gating vector:

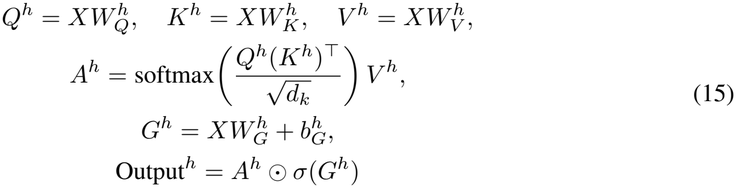

where *h* indexes the attention head, *σ* denotes the Sigmoid function and ⊙ represents element-wise multiplication. This design allows the model to dynamically control information flow at each position.

### B.3 Pretrain Dataset

#### Reference Genome

We use the GRCh38 (hg38) human reference genome as the primary source for pretraining. Since these sequences are directly derived from the reference genome, no additional alignment is required. Each sequence is assigned its corresponding chromosomal coordinates, with the strand fixed to the positive orientation and the CIGAR string set to a full match, and is directly used for alignment-aware training.

#### Multi-species Data

We incorporate multi-species genomic sequences from DNABERT2 [Zhou et al., 2024], capturing diverse evolutionary patterns across different organisms. The sequences are aligned to the human reference genome (hg38) using Minimap2, yielding strand orientation, CIGAR strings, chromosome identifiers, and absolute genomic positions. Among these sequences, 42% are successfully aligned, while the remaining 58% are unalignable. For unalignable sequences, we default to chromosome 0, forward strand orientation, and a CIGAR string representing a full match.

#### 1000 Genomes Project

We leverage phased variant data from the 1000 Genomes Project [Abecasis et al., 2010], which provides high-coverage Illumina sequencing data with phased SNVs and INDELs. Importantly, we utilize the haplotype-resolved information, separately processing variants from paternal (hap0) and maternal (hap1) chromosomes. This enables the model to learn chromosome-specific variation patterns and capture the diploid nature of human genomes. Each variant is represented with its genomic position, sequence, and CIGAR operation type (match, mismatch, insertion, or deletion).

#### Sampling Strategy

The three data sources are combined via dataset concatenation. To balance their contributions during training, sequences from the hg38, multi-species, and 1000 Genomes datasets are sampled at rates of 15%, 45%, and 40% per epoch, respectively. In addition, the training data consist of both contiguous and non-contiguous genomic sequences, where non-contiguous sequences account for 15% of the total samples, while the remaining 85% are contiguous sequences. Furthermore, reverse-complement augmentation is applied with a 30% probability across all datasets to enhance strand-invariant modeling.

For sequences that cannot be reliably aligned, we adopt the default input format described in Appendix I.3.

For pretraining on non-contiguous sequences, we further construct inputs by randomly sampling tokens from random chromosomes and assembling them into non-contiguous sequences, together with their corresponding alignment annotations.

### B.4 Training Details

We pretrain USM-AA-100M and USM-AA-1B using 8 NVIDIA H20 GPUs, respectively. Both models are optimized with AdamW [Loshchilov and Hutter, 2017], using a learning rate of 1 × 10^−4^ and a weight decay of 0.1. A cosine learning rate schedule with 10,000 warmup steps is adopted. Training is conducted for 20 epochs with a fixed token length of 256 and a masking ratio of 15%. USM-AA-100M is trained with a batch size of 240 per GPU, while USM-AA-1B uses a batch size of 160 per GPU. Distributed training is implemented using DeepSpeed [Rajbhandari et al., 2020], with ZeRO Stage-1 optimization, BFloat16 mixed-precision training, and activation checkpointing enabled to improve memory efficiency and scalability. The total pretraining budget is approximately 192 GPU hours for USM-AA-100M (24 hours on 8 GPUs) and 1,536 GPU hours for USM-AA-1B (8 days on 8 GPUs).

### B.5 Alignment Encoding

#### Chromosome Encoding

Chromosome identifiers are encoded as integers using a 1-based indexing scheme, as shown in Table 5. For sequences without valid alignments, a default chromosome ID of 0 is assigned.

**Table 5:** Chromosome encoding scheme used in USM-AA.

| ID | Chromosome | ID | Chromosome |
|---|---|---|---|
| 0 | No alignment | 13 | chr13 |
| 1 | chr1 | 14 | chr14 |
| 2 | chr2 | 15 | chr15 |
| 3 | chr3 | 16 | chr16 |
| 4 | chr4 | 17 | chr17 |
| 5 | chr5 | 18 | chr18 |
| 6 | chr6 | 19 | chr19 |
| 7 | chr7 | 20 | chr20 |
| 8 | chr8 | 21 | chr21 |
| 9 | chr9 | 22 | chr22 |
| 10 | chr10 | 23 | chrX |
| 11 | chr11 | 24 | chrY |
| 12 | chr12 |  |  |

#### Strand Encoding

The strand orientation is encoded as a binary value: 0 for forward strand (+) and 1 for reverse strand (-). During training, reverse-complement augmentation randomly flips sequences to the reverse strand with 30% probability. When a sequence is reverse-complemented, the token sequence is reversed and each nucleotide is replaced with its complement (A↔T, C↔G).

#### CIGAR Encoding

The CIGAR operation encoding scheme follows the SAM/BAM specification, as shown in Table 6.

**Table 6:** CIGAR operation encoding scheme used in USM-AA.

| **ID** | **Operation** | **Symbol** | **Description** |
|---|---|---|---|
| 0 | Match/Mismatch | M | Alignment match (can be match or mismatch) |
| 1 | Insertion | I | Insertion to the reference |
| 2 | Deletion | D | Deletion from the reference |
| 3 | Skip | N | Skipped region from the reference |
| 4 | Soft Clip | S | Soft clipping (sequence present) |
| 5 | Hard Clip | H | Hard clipping (sequence not present) |
| 6 | Padding | P | Padding (silent deletion from padded reference) |
| 7 | Sequence Match | = | Sequence match (exact match) |
| 8 | Sequence Mismatch | X | Sequence mismatch (SNV) |
| 9 | Back | B | Back (reserved) |

#### Position Encoding

Genomic positions are encoded as absolute coordinates on the reference genome (0-based). For the special tokens `<s>` (BOS) and `</s>` (EOS), positions are set to 0 and the chromosome length minus 1, respectively. Each k-mer token is assigned the genomic position of its first nucleotide.

#### Data Augmentation

For sequences with multiple valid alignments, we retain the top 10 alignment candidates ranked by mapping quality. During training, one alignment is randomly sampled from this set for each iteration, which serves as a form of data augmentation. This strategy is consistently applied across all fine-tuning experiments.

### B.6 Dynamic K-mer Tokenizer

We adopt a dynamic *k*-mer tokenization scheme with *k* = 6, resulting in a vocabulary size of 19,552 tokens. Unlike traditional DNA tokenizers that only use the four standard nucleotides (A, C, G, T), our tokenizer includes an additional “N” symbol to represent unknown or ambiguous bases in reference sequences, yielding a 5-letter alphabet.

The “dynamic” mode generates all possible *k*-mers from length 1 to *k*, computed as:

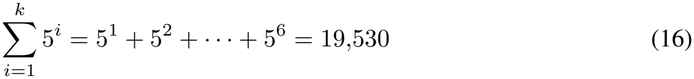

The vocabulary additionally includes 22 special tokens, including the standard tokens `<unk>, <s>, </s>, <pad>`, and `<mask>`, with the remaining tokens reserved for future use.

This yields a total vocabulary size of 19,530 + 22 = 19,552 tokens.

## C Expression Quantitative Trait Loci Prediction

### C.1 Data

The eQTL prediction task is derived from DNALONGBENCH [Cheng et al., 2025] and follows the benchmark defined by Enformer [Avsec et al., 2021]. Statistical fine-mapping variants were collected from GTEx v8 [Consortium, 2020] using SuSiE, a statistical fine-mapping approach widely used to prioritize putative causal variants underlying complex traits [Yuan et al., 2024], and the top nine cell types by variant count were retained. Each sample is a pair of DNA sequences (reference and alternative) spanning the region between the variant and the TSS of the target gene, up to 450 kb, plus an additional 3 kb downstream of the TSS; the two sequences differ only at the variant position. Regions shorter than 450 kb are padded with N bases to a fixed length of 450,000 bp, and other variants inside the region are masked. Positives are variants with posterior inclusion probability (PIP) *>* 0.9, while negatives are matched variants with PIP *<* 0.01 under specified *Z*-score thresholds. The dataset is split into training, validation, and test sets in an 80:10:10 ratio, and comprises 31,282 samples in total. Table 7 lists the cell types and their abbreviations.

**Table 7:** Cell types and abbreviations of the eQTL benchmark. Cell type abbreviations follow DNA-LONGBENCH; Thyroid has no abbreviation.

| **Cell type** | **Abbrev.** |
|---|---|
| Cells – Cultured fibroblasts | CCF |
| Whole Blood | WB |
| Thyroid | – |
| Skin – Not Sun Exposed (Suprapubic) | SNSES |
| Skin – Sun Exposed (Lower Leg) | SSELL |
| Muscle – Skeletal | MS |
| Nerve – Tibial | NT |
| Artery – Tibial | AT |
| Adipose – Subcutaneous | AS |

### C.2 Input Design

The original protocol pads every sample into a 450,000 bp contiguous sequence carrying a single discriminating nucleotide. We instead feed USM-AA only the two informative segments, namely a window centered on the variant and a window centered on the TSS, and let AARoPE encode their absolute genomic coordinates so that the model directly perceives the true variant–TSS distance without consuming the intervening region. Because the reference and alternative inputs differ only at the variant token, the comparison reduces to comparing the two conditions within one coordinate-aware representation rather than processing a full ultra-long sequence.

### C.3 Implementation Details

USM-AA-100M and USM-AA-1B are fine-tuned end-to-end with an average pooling classification head under full-parameter fine-tuning, using the discrete-segment input design described above. Each model is trained on 8 NVIDIA H20 GPUs with the FusedAdam optimizer (learning rate 5 × 10^−5^, weight decay 0.1), a cosine learning rate schedule with 100 warmup steps, 10 epochs, a maximum sequence length of 2,048 tokens, and a batch size of 1 per GPU. Following the original benchmark, we report AUROC for each cell type and the average across the nine cell types. The same fine-tuning setup is used for the ETGP task in Appendix D.

### C.4 Results

Table 8 reports the per-cell-type results. Baseline results are cited from the original DNALONG-BENCH study for fair comparison, and USM-AA is fine-tuned on the same dataset splits. Both USM-AA scales rank first or second in every cell type and clearly exceed the Expert Model and all DNA foundation model baselines, whose performance is far from satisfactory in most cell types.

**Table 8:**
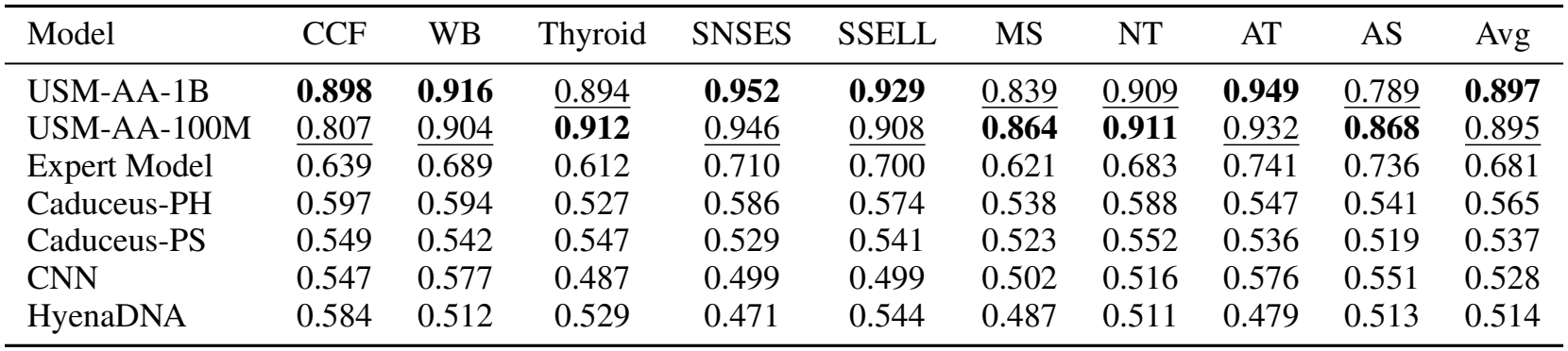
Expression quantitative trait loci prediction across nine human cell types. Values are AUROC. Best results are shown in **bold**, second-best results are underlined. Cell type abbreviations follow Table 7.

Table 9 ablates the alignment-aware components of USM-AA-100M on the eQTL task. Removing the alignment embeddings (w/o AEM) causes a moderate drop in average AUROC, while replacing AARoPE with relative positional encoding (w/o AARoPE) leads to a much larger degradation of 0.066, indicating that the coordinate-aware positional information, rather than the alignment embeddings alone, is the primary source of the variant-comparison ability. Removing both components yields the worst average AUROC of 0.822, slightly below removing AARoPE alone, so the two components contribute largely through the same pathway, with only a small complementary effect.

**Table 9:**
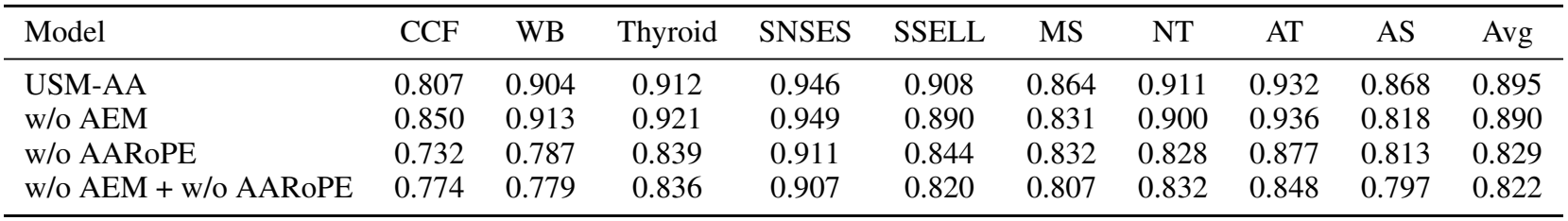
Ablation of alignment-aware components on the eQTL task (USM-AA-100M). Values are AUROC. “w/o AARoPE” uses relative positional encoding instead of AARoPE; “w/o AEM” removes the alignment embeddings. Cell type abbreviations follow Table 7.

| Model | CCF | WB | Thyroid | SNSES | SSELL | MS | NT | AT | AS | Avg |
|---|---|---|---|---|---|---|---|---|---|---|
| USM-AA | 0.807 | 0.904 | 0.912 | 0.946 | 0.908 | 0.864 | 0.911 | 0.932 | 0.868 | 0.895 |
| w/o AEM | 0.850 | 0.913 | 0.921 | 0.949 | 0.890 | 0.831 | 0.900 | 0.936 | 0.818 | 0.890 |
| w/o AARoPE | 0.732 | 0.787 | 0.839 | 0.911 | 0.844 | 0.832 | 0.828 | 0.877 | 0.813 | 0.829 |
| w/o AEM + w/o AARoPE | 0.774 | 0.779 | 0.836 | 0.907 | 0.820 | 0.807 | 0.832 | 0.848 | 0.797 | 0.822 |

## D Enhancer Target Gene Prediction

### D.1 Data

The ETGP task is derived from DNALONGBENCH [Cheng et al., 2025]. The dataset contains 2,602 experimentally verified enhancer-promoter interactions in K562 cells from CRISPRi-based studies [Fulco et al., 2019, Gasperini et al., 2019, Schraivogel et al., 2020], split into 2,066 training, 266 validation, and 270 test samples. Only enhancer-gene pairs within 450 kb of the gene TSS are retained, making this a challenging long-range genomic interaction prediction problem. We compare against the official DNALONGBENCH baselines; baseline results are cited from the original study for fair comparison.

### D.2 Input Design

The original DNALONGBENCH protocol extracts sequences from 500 bp upstream of the enhancer to 3 kb downstream of the TSS, pads them with N bases to 450 kb, and masks other enhancers within the region. As in eQTL, we instead retain only the two informative segments: the enhancer region (extended by 500 bp on each side) and a 6,000 bp window centered on the TSS, with AARoPE encoding their absolute genomic coordinates so that the enhancer–promoter interaction can be modeled without consuming the intervening sequence. This substantially reduces the computational cost of ultra-long inputs while preserving the essential positional information. The fine-tuning setup follows Appendix C.

## E TCGA PanCanAtlas Cancer Classification

### E.1 Data

The TCGA Pan-Cancer Atlas [Hoadley et al., 2018] profiles 33 cancer types across multiple molecular layers and provides a comprehensive resource for integrative cancer analysis. The dataset is publicly available at https://gdc.cancer.gov/about-data/publications/pancanatlas.

Somatic variants were parsed from MAF files, with each record containing chromosomal position, reference and alternate alleles, and variant type. Samples with fewer than 10 mutations were excluded. The remaining samples were stratified by cancer type and randomly split into training (60%), validation (20%), and test (20%) sets.

The resulting dataset consists of 9,824 samples spanning all 33 cancer types. Each sample is represented as a set of non-contiguous somatic variants, including SNVs and INDELs, reflecting the highly heterogeneous mutational landscapes characteristic of human cancers. Detailed statistics on mutation counts and variant composition are summarized in Table 10.

**Table 10:** Statistical Summary of Genomic Variants in TCGA Pan-Cancer Atlas Dataset.

| **Dataset** | **Samples** | **SNV Range** | **SNV Median** | **INDEL Range** | **INDEL Median** |
|---|---|---|---|---|---|
| Training | 5,880 | 6–55,145 | 90 | 0–2,539 | 4 |
| Validation | 1,952 | 8–48,012 | 92 | 0–2,270 | 5 |
| Test | 1,992 | 7–39,582 | 91 | 0–1,951 | 4 |
| Total | 9,824 | 6–55,145 | 91 | 0–2,539 | 4 |

For traditional machine learning baselines (XGBoost and Logistic Regression), we perform feature extraction by computing the counts of SNVs, INDELs, and multi-nucleotide variants for each chromosome as well as their genome-wide totals. For SNVs, we further quantify the number of variants corresponding to each nucleotide substitution (A, T, C, and G). These features are then used as inputs for classification.

### E.2 Implementation Details

Since existing GFMs are not designed for non-contiguous mutation inputs, we compare our approach with CNN and Transformer baselines. The Transformer model uses the USM-AA architecture trained from scratch. The CNN baseline adopts a 12-layer residual convolutional architecture with multi-scale kernels and the same embedding dimension (768) as USM-AA-100M.

All models are trained on 8 NVIDIA H20 GPUs using the AdamW optimizer with a weight decay of 0.1 and a cosine learning rate schedule with 100 warmup steps. For USM-AA-100M, USM-AA-1B, and the Transformer trained from scratch, we use a batch size of 1 per GPU and a learning rate of 5 × 10^−5^. The CNN baseline uses a batch size of 8 per GPU and a learning rate of 1 × 10^−4^. All models are trained for 20 epochs with a maximum sequence length of 65,536 tokens; due to GPU memory constraints, the sequence length for USM-AA-1B is limited to 20,480 tokens. Full results are reported in Table 2.

## F NT Benchmarks

### F.1 Data

The NT benchmark consists of 18 downstream tasks designed to provide a unified evaluation framework for genomic sequence modeling. These datasets are available for down-load at https://huggingface.co/datasets/InstaDeepAI/nucleotide_transformer_downstream_tasks_revised.

These tasks cover both binary and multi-class classification settings and are exclusively derived from human genomic data. They are grouped into three biologically relevant categories: Chromatin Profiles, Regulatory Elements, and Splicing-related tasks. Together, these tasks span a wide range of functional genomics prediction problems.

Detailed statistics of the benchmark, including the number of training and test sequences, label cardinality, and sequence length for each task, are summarized in Table 11.

**Table 11:** Composition of NT Datasets.

| **Dataset** | **Category** | **Num. Sequences** | **Num. Classes** | **Sequence Length** |
|---|---|---|---|---|
| Promoter all | Regulatory Elements | 31,584 | 2 | 300 |
| Promoter tata | Regulatory Elements | 5,274 | 2 | 300 |
| Promoter no tata | Regulatory Elements | 31,372 | 2 | 300 |
| Enhancers | Regulatory Elements | 33,000 | 2 | 400 |
| Enhancers types | Regulatory Elements | 33,000 | 2 | 400 |
| Splice sites all | Splicing | 33,000 | 2 | 600 |
| Splice sites acceptor | Splicing | 33,000 | 2 | 600 |
| Splice sites donor | Splicing | 33,000 | 2 | 600 |
| H2AFZ | Chromatin Profiles | 33,000 | 2 | 1,000 |
| H3K27ac | Chromatin Profiles | 31,616 | 2 | 1,000 |
| H3K27me3 | Chromatin Profiles | 33,000 | 2 | 1,000 |
| H3K36me3 | Chromatin Profiles | 33,000 | 2 | 1,000 |
| H3K4me1 | Chromatin Profiles | 33,000 | 2 | 1,000 |
| H3K4me2 | Chromatin Profiles | 32,138 | 2 | 1,000 |
| H3K4me3 | Chromatin Profiles | 30,776 | 2 | 1,000 |
| H3K9ac | Chromatin Profiles | 24,278 | 2 | 1,000 |
| H3K9me3 | Chromatin Profiles | 28,288 | 2 | 1,000 |
| H4K20me1 | Chromatin Profiles | 32,270 | 2 | 1,000 |

### F.2 Implementation Details

For fine-tuning on the Nucleotide Transformer benchmark tasks, we used pretrained USM-AA as the backbone and attached an average pooling classification head, with all parameters fine-tuned. All experiments were conducted on 4 NVIDIA A100 GPUs, and the detailed hyperparameters are summarized in Table 12.

**Table 12:** Hyperparameters for fine-tuning on Nucleotide Transformer benchmark tasks.

| **Hyperparameter** | **USM-AA-100M** | **USM-AA-1B** |
|---|---|---|
| Optimizer | AdamW | AdamW |
| Learning rate | 5 × 10<sup>−5</sup> | 5 × 10<sup>−5</sup> |
| Weight decay | 0.1 | 0.1 |
| Batch size per GPU | 32 | 16 |
| Number of GPUs | 4 | 4 |
| Max sequence length | 1024 | 1024 |
| Warmup steps | 100 | 100 |
| LR scheduler | Cosine | Cosine |
| Dropout | 0.1 | 0.1 |
| Gradient clipping | 1.0 | 1.0 |
| Precision | BFloat16 | BFloat16 |

For model selection, we employed early stopping based on validation performance, and reported final results on the test set. We performed 10-fold cross-validation using random seeds from 0 to 9, reporting results as the mean ± standard deviation across the folds. For the Chromatin Profiles and enhancer tasks, models were trained for 10 epochs, while for all other tasks, training was conducted for 25 epochs.

### F.3 NT Results

The detailed results are in Table 13. All models are evaluated using 10-fold cross-validation with random seeds from 0 to 9, applying early stopping based on validation MCC. The reported results of the baseline methods (DNABERT2, NT, HyenaDNA, Enformer, and SPACE) are directly taken from the original NT and SPACE studies. For the two NTv3-650M variants (pre and post), we fine-tuned the official checkpoints using the same hyperparameters and evaluation protocol as USM-AA, ensuring a fair comparison.

**Table 13:** Results of Nucleotide Transformer downstream tasks. This benchmark includes three categories (18 tasks) derived from human samples. Values are reported as mean ± standard deviation. Best results are shown in **bold**, and second-best results are underlined. Baseline results were sourced from SPACE and NT, except for the two NTv3 variants, which were fine-tuned by us using the same hyperparameters as USM-AA.

| Model | Params | Chromatin Profiles |  |  |  |  |  |
|---|---|---|---|---|---|---|---|
|  |  | H2AFZ | H3K27ac | H3K27me3 | H3K36me3 | H3K4me1 | H3K4me2 |
| DNABERT2 | 117M | 0.490 ± 0.013 | 0.491 ± 0.010 | 0.599 ± 0.010 | 0.637 ± 0.007 | 0.490 ± 0.008 | 0.558 ± 0.013 |
| HyenaDNA-32KB | 3.9M | 0.467 ± 0.012 | 0.421 ± 0.010 | 0.550 ± 0.009 | 0.553 ± 0.011 | 0.423 ± 0.016 | 0.515 ± 0.018 |
| NT-HumanRef | 500M | 0.465 ± 0.011 | 0.457 ± 0.010 | 0.589 ± 0.009 | 0.594 ± 0.004 | 0.468 ± 0.007 | 0.527 ± 0.011 |
| NT-Multispecies | 2.5B | 0.503 ± 0.010 | 0.481 ± 0.020 | 0.593 ± 0.016 | 0.635 ± 0.016 | 0.481 ± 0.012 | 0.552 ± 0.022 |
| NTv3-pre | 650M | 0.463 ± 0.015 | 0.483 ± 0.026 | 0.569 ± 0.025 | 0.597 ± 0.006 | 0.472 ± 0.027 | 0.515 ± 0.018 |
| NTv3-post | 650M | 0.496 ± 0.017 | 0.595 ± 0.030 | 0.605 ± 0.016 | 0.643 ± 0.012 | 0.559 ± 0.022 | 0.622 ± 0.018 |
| Enformer | 252M | 0.522 ± 0.019 | 0.520 ± 0.015 | 0.552 ± 0.007 | 0.567 ± 0.017 | 0.504 ± 0.021 | 0.626 ± 0.015 |
| SPACE | 588M | 0.548 ± 0.005 | 0.547 ± 0.007 | 0.586 ± 0.010 | 0.602 ± 0.005 | 0.543 ± 0.009 | 0.640 ± 0.007 |
| USM-AA | 122M | 0.584 ± 0.029 | 0.592 ± 0.016 | 0.625 ± 0.035 | 0.675 ± 0.024 | 0.552 ± 0.038 | 0.640 ± 0.029 |
| USM-AA | 1.16B | **0.633 ± 0.036** | **0.603 ± 0.018** | **0.647 ± 0.019** | **0.686 ± 0.009** | **0.598 ± 0.049** | **0.699 ± 0.037** |
| Model | Params | Chromatin Profiles |  |  |  | Regulatory Elements |  |
|  |  | H3K4me3 | H3K9ac | H3K9me3 | H4K20me1 | Enhancers | Enhancers(types) |
| DNABERT2 | 117M | 0.646 ± 0.008 | 0.564 ± 0.013 | 0.443 ± 0.025 | 0.655 ± 0.011 | 0.517 ± 0.011 | 0.476 ± 0.009 |
| HyenaDNA-32KB | 3.9M | 0.603 ± 0.020 | 0.487 ± 0.025 | 0.419 ± 0.030 | 0.590 ± 0.007 | 0.476 ± 0.021 | 0.445 ± 0.009 |
| NT-HumanRef | 500M | 0.622 ± 0.013 | 0.524 ± 0.013 | 0.433 ± 0.009 | 0.634 ± 0.013 | 0.515 ± 0.019 | 0.477 ± 0.014 |
| NT-Multispecies | 2.5B | 0.618 ± 0.015 | 0.527 ± 0.017 | 0.447 ± 0.018 | 0.650 ± 0.014 | 0.527 ± 0.012 | 0.484 ± 0.012 |
| NTv3-pre | 650M | 0.645 ± 0.030 | 0.639 ± 0.035 | 0.492 ± 0.042 | 0.622 ± 0.018 | 0.557 ± 0.030 | 0.509 ± 0.018 |
| NTv3-post | 650M | **0.693 ± 0.023** | **0.697 ± 0.025** | 0.515 ± 0.040 | 0.664 ± 0.014 | 0.658 ± 0.013 | **0.622 ± 0.019** |
| Enformer | 252M | 0.635 ± 0.019 | 0.593 ± 0.020 | 0.453 ± 0.016 | 0.606 ± 0.016 | 0.614 ± 0.010 | 0.573 ± 0.013 |
| SPACE | 588M | 0.661 ± 0.025 | 0.635 ± 0.016 | 0.490 ± 0.011 | 0.650 ± 0.011 | 0.631 ± 0.007 | 0.583 ± 0.008 |
| USM-AA | 122M | 0.676 ± 0.025 | 0.639 ± 0.016 | 0.530 ± 0.057 | 0.703 ± 0.020 | 0.623 ± 0.034 | 0.609 ± 0.019 |
| USM-AA | 1.16B | 0.651 ± 0.020 | 0.632 ± 0.015 | **0.682 ± 0.037** | **0.725 ± 0.013** | **0.672 ± 0.031** | 0.610 ± 0.032 |
| Model | Params | Regulatory Elements |  |  | Splicing |  |  |
|  |  | All | NoTATA | TATA | Donors | Acceptors | All |
| DNABERT2 | 117M | 0.754 ± 0.009 | 0.769 ± 0.009 | 0.784 ± 0.036 | 0.837 ± 0.006 | 0.855 ± 0.005 | 0.861 ± 0.004 |
| HyenaDNA-32KB | 3.9M | 0.698 ± 0.011 | 0.729 ± 0.009 | 0.666 ± 0.041 | 0.808 ± 0.009 | 0.907 ± 0.018 | 0.915 ± 0.047 |
| NT-HumanRef | 500M | 0.734 ± 0.013 | 0.738 ± 0.008 | 0.831 ± 0.022 | 0.941 ± 0.004 | 0.939 ± 0.003 | 0.952 ± 0.003 |
| NT-Multispecies | 2.5B | 0.761 ± 0.009 | 0.773 ± 0.010 | **0.944 ± 0.016** | **0.958 ± 0.003** | **0.964 ± 0.003** | **0.970 ± 0.002** |
| NTv3-pre | 650M | 0.787 ± 0.017 | 0.803 ± 0.015 | 0.900 ± 0.039 | 0.952 ± 0.013 | 0.935 ± 0.010 | 0.921 ± 0.011 |
| NTv3-post | 650M | 0.774 ± 0.013 | 0.791 ± 0.015 | 0.875 ± 0.036 | 0.850 ± 0.009 | 0.848 ± 0.011 | 0.864 ± 0.012 |
| Enformer | 252M | 0.745 ± 0.012 | 0.763 ± 0.012 | 0.793 ± 0.026 | 0.749 ± 0.007 | 0.739 ± 0.011 | 0.780 ± 0.007 |
| SPACE | 588M | 0.764 ± 0.012 | 0.776 ± 0.011 | 0.838 ± 0.028 | 0.942 ± 0.006 | 0.902 ± 0.004 | 0.906 ± 0.003 |
| USM-AA | 122M | **0.817 ± 0.018** | **0.820 ± 0.019** | 0.858 ± 0.018 | 0.937 ± 0.011 | 0.935 ± 0.007 | 0.954 ± 0.005 |
| USM-AA | 1.16B | 0.802 ± 0.015 | 0.806 ± 0.007 | 0.866 ± 0.028 | 0.954 ± 0.004 | 0.943 ± 0.005 | 0.957 ± 0.005 |

## G GUE Benchmarks

### G.1 Overview

The GUE benchmark [Zhou et al., 2024] is a standardized evaluation suite designed to systematically assess the sequence-level understanding capability of genome foundation models across a diverse set of biologically meaningful tasks. We evaluate USM-AA on the GUE benchmark to assess its cross-species generalization ability, which includes tasks such as yeast epigenetic mark prediction, human and mouse transcription factor (TF) binding prediction, and COVID variant classification (CVC).

### G.2 Data

GUE focuses on short to medium length genomic sequences, covering a broad range of functional genomics problems, including promoter detection, transcription factor binding prediction, splice site recognition, epigenetic mark prediction, and viral variant classification. The bench-mark spans multiple species including human, mouse, yeast, and viruses, comprising a total of 28 datasets across seven task categories. These datasets are available for download at https://github.com/MAGICS-LAB/DNABERT_2. The composition of the benchmark, including the species, task categories, number of datasets, number of classes, and sequence lengths, is summarized in Table 14.

**Table 14:** The Composition of GUE Datasets.

| Species | Task | Num. Datasets | Num. Classes | Sequence Length |
|---|---|---|---|---|
| Human | Core Promoter Detection (CPD) | 3 | 2 | 70 |
|  | Transcription Factor Prediction (TF-H) | 5 | 2 | 100 |
|  | Promoter Detection (PD) | 3 | 2 | 300 |
|  | Splice Site Detection (SSD) | 1 | 3 | 400 |
| Mouse | Transcription Factor Prediction (TF-M) | 5 | 2 | 100 |
| Yeast | Epigenetic Marks Prediction (EMP) | 10 | 2 | 500 |
| Virus | Covid Variant Classification (CVC) | 1 | 9 | 1000 |

### G.3 Implementation Details

For fine-tuning on the GUE benchmark tasks, we used pretrained USM-AA as the backbone and attached an average pooling classification head, with all parameters fine-tuned. All experiments are conducted using 4 NVIDIA A100 GPUs, and the detailed hyperparameters are summarized in Table 15.

**Table 15:** Hyperparameters for fine-tuning on GUE benchmark tasks.

| **Hyperparameter** | **USM-AA-100M** | **USM-AA-1B** |
|---|---|---|
| Optimizer | AdamW | AdamW |
| Learning rate | 5 × 10<sup>−5</sup> | 5 × 10<sup>−5</sup> |
| Weight decay | 0.1 | 0.1 |
| Batch size per GPU | 32 | 16 |
| Max sequence length | 512 | 512 |
| Warmup steps | 100 | 100 |
| LR scheduler | Cosine | Cosine |
| Dropout | 0.1 | 0.1 |
| Gradient clipping | 1.0 | 1.0 |
| Precision | BFloat16 | BFloat16 |

For model selection, we employed early stopping based on validation performance. All tasks were trained for 10 epochs, except for the CVC task, which was trained for 20 epochs due to slower convergence and used a larger batch size of 64.

### G.4 Results

Following prior work, F1-score is used for the COVID Variant Classification (CVC) task, while MCC is used for all other tasks. USM-AA is compared with DNABERT2, NT-Multispecies, Enformer [Avsec et al., 2021], and SPACE, using baseline results from the original studies for fair comparison.

As shown in Table 16, USM-AA attains the highest performance on four of the seven major task categories, with detailed results across all 28 datasets. In contrast, the comparatively lower performance on TF-M and TF-H prediction tasks in human and mouse is likely due to our model being pretrained without functional track data compared with SPACE.

**Table 16:** Results on GUE benchmark (28 tasks). Best results are in **bold**, second best are underlined. Baseline results were sourced from SPACE and DNABERT2.

| Model | Params | EMP |  |  |  |  |  |  |  |  |  |
|---|---|---|---|---|---|---|---|---|---|---|---|
|  |  | H3 | H3K14ac | H3K36me3 | H3K4me1 | H3K4me2 | H3K4me3 | H3K79me3 | H3K9ac |  |  |
| DNABERT2 | 117M | 78.27 | 52.57 | 56.88 | 50.52 | 31.13 | 36.27 | 67.39 | 55.63 |  |  |
| NT-Multispecies | 2.5B | 78.77 | 56.20 | 61.99 | 55.30 | 36.49 | 40.34 | 64.70 | 56.01 |  |  |
| Enformer | 252M | 70.65 | 37.87 | 42.41 | 34.00 | 29.65 | 22.19 | 55.69 | 49.35 |  |  |
| SPACE | 588M | **79.53** | 54.12 | 54.82 | 50.92 | **43.80** | **49.47** | 66.93 | 59.29 |  |  |
| USM-AA | 122M | 74.41 | 55.46 | 62.71 | **55.70** | 35.28 | 43.43 | 66.67 | 56.74 |  |  |
| USM-AA | 1.16B | 78.18 | **58.48** | **63.21** | 53.63 | 33.74 | 48.41 | **69.62** | **62.59** |  |  |
| Model | Params | EMP |  | PD |  |  | CPD |  |  | CVC | Splice |
|  |  | H4 | H4ac | all | notata | tata | all | notata | tata | Virus | Splice |
| DNABERT2 | 117M | 80.71 | 50.43 | 86.77 | 94.27 | 71.59 | 69.37 | 68.04 | 74.17 | 71.02 | 84.99 |
| NT-Multispecies | 2.5B | 81.67 | 49.13 | 91.01 | 94.00 | **79.43** | 70.33 | 71.58 | 72.97 | **73.04** | 89.35 |
| Enformer | 252M | 76.32 | 32.90 | 85.68 | 92.92 | 69.63 | 60.94 | 66.46 | 46.21 | 61.33 | 81.55 |
| SPACE | 588M | 81.25 | 53.09 | 91.90 | 94.23 | 79.13 | 68.18 | 68.04 | **79.23** | 70.26 | 87.48 |
| USM-AA | 122M | 80.87 | 51.63 | 95.71 | 100.00 | 74.66 | **96.03** | 99.74 | 71.43 | 62.62 | 89.58 |
| USM-AA | 1.16B | **83.80** | **57.38** | **95.93** | **100.00** | 75.37 | 95.61 | **99.74** | 74.22 | 69.15 | **91.57** |
| Model | Params | TF-H |  |  |  |  | TF-M |  |  |  |  |
|  |  | 0 | 1 | 2 | 3 | 4 | 0 | 1 | 2 | 3 | 4 |
| DNABERT2 | 117M | **71.99** | 76.06 | 66.52 | 58.54 | 77.43 | 56.76 | 84.77 | 79.32 | 66.47 | **52.66** |
| NT-Multispecies | 2.5B | 66.64 | 70.28 | 58.72 | 51.65 | 69.34 | 63.31 | 83.76 | 71.52 | 69.44 | 47.07 |
| Enformer | 252M | 69.42 | 72.76 | **77.88** | **66.41** | 81.89 | **67.15** | 81.56 | 85.99 | 67.88 | 44.03 |
| SPACE | 588M | 69.02 | **76.49** | 76.45 | 66.08 | **82.91** | 65.94 | 84.91 | **90.30** | **86.72** | 50.66 |
| USM-AA | 122M | 65.55 | 72.15 | 65.08 | 62.80 | 72.21 | 66.05 | **86.04** | 82.33 | 77.52 | 49.81 |
| USM-AA | 1.16B | 65.80 | 71.06 | 62.24 | 56.44 | 75.43 | 63.31 | 84.88 | 81.80 | 85.91 | 50.83 |

## H Genomic Benchmarks

### H.1 Overview

To evaluate USM-AA across diverse genomic classification tasks, we benchmarked it on the Genomic Benchmarks (GB) dataset [Grešová et al., 2023]. This benchmark encompasses enhancer detection, coding versus intergenic region classification, species discrimination, regulatory element prediction, non-TATA promoter identification, and open chromatin region classification across multiple organisms.

We compared USM-AA with baseline models, including HyenaDNA, Caduceus [Schiff et al., 2024], Enformer, and SPACE. All experiments were conducted following the same protocol as SPACE, while results for Caduceus and SPACE were directly adopted from their original publications to ensure a fair comparison. For the Drosophila dataset, since Caduceus did not report results, we only included SPACE and Enformer as baselines.

### H.2 Data

The Genomic Benchmarks provide a compact evaluation framework for regulatory sequence modeling, consisting of nine datasets that reflect key regulatory phenomena, including transcription initiation, distal regulation, and chromatin openness. The benchmark spans genomic data from human, mouse, drosophila and nematoda. An overview of the individual datasets is provided in Table 17. These datasets are available for download at https://github.com/ML-Bioinfo-CEITEC/genomic_benchmarks.

**Table 17:** Composition of Genomic Benchmarks.

| **Dataset** | **Num. Sequences** | **Num. Classes** | **Mean Seq. Length** |
|---|---|---|---|
| Dummy mouse enhancers ensembl | 1,210 | 2 | 2,369.58 |
| Demo coding vs intergenomic seqs | 100,000 | 2 | 200.00 |
| Demo human or worm | 100,000 | 2 | 200.00 |
| Drosophila enhancers stark | 6,914 | 2 | 2,118.12 |
| Human enhancers cohn | 27,791 | 2 | 500.00 |
| Human enhancers ensembl | 154,842 | 2 | 268.86 |
| Human ensembl regulatory | 289,061 | 2 | 429.92 |
| Human nontata promoters | 36,131 | 2 | 251.00 |
| Human ocr ensembl | 174,756 | 2 | 326.35 |

### H.3 Implementation Details

For fine-tuning on Genomic Benchmarks tasks, we used pretrained USM-AA as the backbone and attached an average pooling classification head, with all parameters fine-tuned. All experiments were conducted on 4 NVIDIA A100 GPUs. The detailed hyperparameters are summarized in Table 18.

**Table 18:** Hyperparameters for fine-tuning on Genomic Benchmarks tasks.

| **Hyperparameter** | **USM-AA-100M** | **USM-AA-1B** |
|---|---|---|
| Optimizer | AdamW | AdamW |
| Learning rate | 5 × 10<sup>−5</sup> | 5 × 10<sup>−5</sup> |
| Weight decay | 0.1 | 0.1 |
| Batch size per GPU | 32 | 16 |
| Max sequence length | 4096 | 4096 |
| Warmup steps | 100 | 100 |
| LR scheduler | Cosine | Cosine |
| Dropout | 0.1 | 0.1 |
| Gradient clipping | 1.0 | 1.0 |
| Precision | BFloat16 | BFloat16 |

For model selection, we performed cross-validation using random seeds 0–4, employing early stopping based on validation performance, consistent with prior work. Note that for the Dummy Mouse Enhancers Ensembl task, the batch size per GPU was set to 4 and the model was trained for 30 epochs due to the small size of the dataset.

### H.4 Results

As shown in Table 19, USM-AA achieves the best performance on six out of nine tasks. Notably, USM-AA demonstrates particularly strong results on the Demo Coding VS. Intergenic and Human VS. Worm classification tasks, achieving near-perfect accuracy. On the Human VS. Worm task, USM-AA-1B achieves an accuracy of 0.999, substantially outperforming all baseline models.

**Table 19:** Results of Genomic Benchmarks. Best scores are **bold**, second-best scores are underlined. The baseline results were sourced from SPACE and Caduceus [Schiff et al., 2024]. Values are reported as mean ± standard deviation.

| Model | Params | Mouse | Demo |  | Drosophila |  |
|---|---|---|---|---|---|---|
|  |  | **Enhancers** | **Coding VS. Interg.** | **Human VS. Worm** | **Enhancers** |  |
| HyenaDNA | 1.6M | 0.780 ± 0.025 | 0.904 ± 0.005 | 0.964 ± 0.002 | – |  |
| Caduceus-PH | 1.9M | 0.754 ± 0.074 | 0.915 ± 0.003 | 0.973 ± 0.001 | – |  |
| Caduceus-PS | 1.9M | 0.793 ± 0.058 | 0.910 ± 0.003 | 0.968 ± 0.002 | – |  |
| Enformer | 252M | 0.835 ± 0.012 | 0.913 ± 0.001 | 0.958 ± 0.001 | 0.613 ± 0.005 |  |
| SPACE | 588M | **0.905 ± 0.010** | 0.922 ± 0.001 | 0.967 ± 0.004 | 0.721 ± 0.016 |  |
| USM-AA | 122M | 0.821 ± 0.016 | 0.976 ± 0.001 | **0.999 ± 0.000** | 0.809 ± 0.021 |  |
| USM-AA | 1.16B | 0.832 ± 0.010 | **0.978 ± 0.001** | **0.999 ± 0.000** | **0.823 ± 0.009** |  |
| Model | Params | Human |  |  |  |  |
|  |  | **Enh. Cohn** | **Enh. Ensembl** | **Regulatory** | **OCR Ensembl** | **Nontata Prom.** |
| HyenaDNA | 1.6M | 0.729 ± 0.014 | 0.849 ± 0.006 | 0.869 ± 0.012 | 0.783 ± 0.007 | 0.944 ± 0.002 |
| Caduceus-PH | 1.9M | 0.747 ± 0.004 | 0.893 ± 0.008 | 0.872 ± 0.011 | 0.828 ± 0.006 | 0.946 ± 0.007 |
| Caduceus-PS | 1.9M | 0.745 ± 0.007 | 0.900 ± 0.006 | 0.873 ± 0.007 | 0.818 ± 0.006 | 0.945 ± 0.010 |
| Enformer | 252M | 0.723 ± 0.001 | 0.844 ± 0.001 | 0.903 ± 0.001 | **0.876 ± 0.001** | 0.878 ± 0.002 |
| SPACE | 588M | **0.769 ± 0.006** | 0.919 ± 0.014 | 0.944 ± 0.002 | 0.854 ± 0.001 | 0.940 ± 0.002 |
| USM-AA | 122M | 0.767 ± 0.009 | 0.910 ± 0.009 | 0.943 ± 0.006 | 0.776 ± 0.011 | 0.986 ± 0.003 |
| USM-AA | 1.16B | 0.768 ± 0.005 | **0.920 ± 0.007** | **0.951 ± 0.004** | 0.792 ± 0.007 | **0.993 ± 0.001** |

For human-specific tasks, USM-AA attains state-of-the-art performance on Enhancers Ensembl, Regulatory element prediction, and Non-TATA promoter identification. The Non-TATA promoter task shows particularly impressive results, with USM-AA-1B achieving 0.993 accuracy. However, on the Mouse Enhancers task, SPACE maintains the best performance, likely due to its incorporation of functional track data during pretraining.

The Drosophila Enhancers task demonstrates USM-AA’s cross-species generalization capability, where it significantly outperforms both Enformer and SPACE despite being pretrained primarily on human genomic data aligned to the hg38 reference.

## I Ablation

### I.1 Alignment-Aware Learning

To assess whether the model can encode positional information through genomic coordinate systems rather than relying on local sequence context, we evaluate the USM-AA-100M model under varying masking ratios during pretraining. When the masking ratio is set to 1.0, all input tokens are masked, forcing the model to rely exclusively on chromosomal coordinate information for token prediction.

Training data are constructed from the hg38 reference genome, comprising 50% contiguous genomic segments and 50% discretely sampled non-contiguous sequences. Non-contiguous sequences are randomly sampled from each chromosome to disrupt local continuity and remove contextual cues. This setup prevents the model from exploiting neighboring token dependencies and encourages reliance on coordinate-based positional encoding, providing a controlled evaluation of alignment-aware representation learning.

As shown in Fig. 4(a), the training dynamics exhibit limited sensitivity to the masking ratio. Across a wide range of masking ratios (0.1 to 0.7), the training loss curves remain largely consistent. Even under full masking (mask ratio = 1.0), where no sequence tokens are observable, the model demon-strates stable convergence, indicating that it can memorize and leverage genomic coordinate information to recover nucleotide identities.

**Figure 4:**
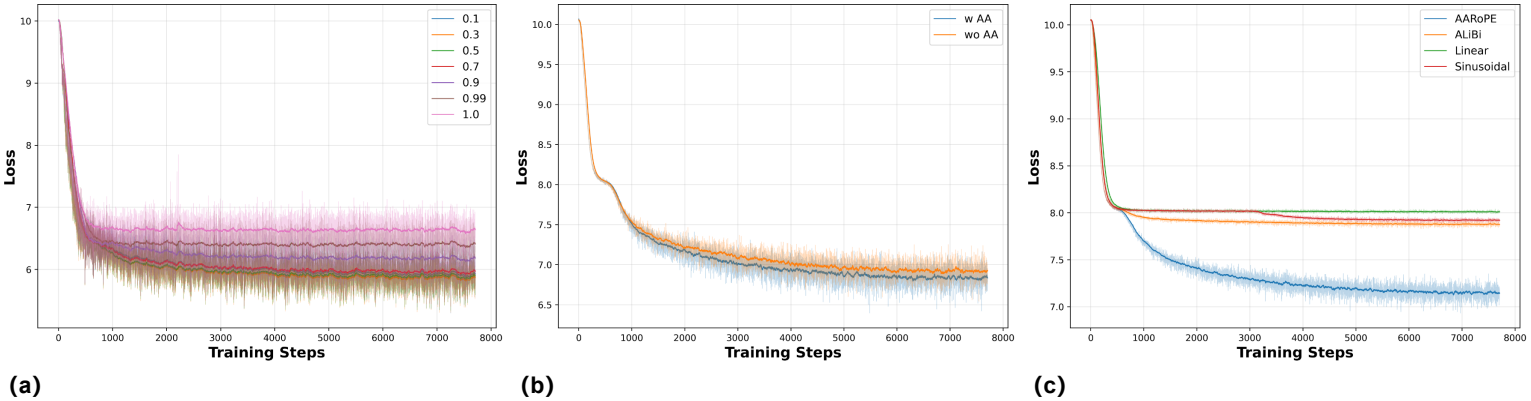
Ablation studies on alignment-aware learning. (a) Training loss curves under different masking ratios. (b) Training loss curves for pretraining with alignment-aware sequences and without alignment information. (c) Training loss curves for different positional encoding methods on the reference genome pretraining task. We compare four approaches: AARoPE, ALiBi, Linear positional encoding, and Sinusoidal positional encoding. All models use the USM-AA-100M architecture with identical hyperparameters. AARoPE demonstrated the lowest loss during pretraining on hybrid pretrain datasets.

To further evaluate the effect of alignment-aware inputs during pretraining, we compare two settings: pretraining on alignment-aware contiguous sequences and on unaligned contiguous sequences. The latter removes explicit coordinate information while preserving sequence continuity, isolating the contribution of alignment signals. As shown in Fig. 4(b), the USM-AA-100M model pretrained with alignment-aware inputs (w/ AA) converges faster than the model pretrained without alignment information (w/o AA), suggesting that incorporating genomic alignment information facilitates more efficient representation learning.

### I.2 Positional Encoding Evaluation

To select an appropriate positional encoding method capable of handling absolute chromosomal coordinates, we evaluated four positional encoding approaches: Sinusoidal positional encoding, AL-iBi, Linear positional encoding, and AARoPE. All experiments were conducted on the reference genome with identical hyperparameters to ensure fair comparison.

#### Experimental Setup

We used the following configuration: batch size of 320, sequence length of 256 tokens, mask ratio of 15%, learning rate of 1×10^−4^ with cosine decay schedule including 10,000 warmup steps, AdamW optimizer with weight decay of 0.1, gradient clipping of 0.5, and training for 10 epochs using bfloat16 precision. The model architecture is based on USM-AA-100M.

#### Data Sampling Strategy

To handle the challenge of modeling long-range genomic dependencies while maintaining computational efficiency, we employed the sparse sampling strategy described in Appendix I.4: given an original contiguous sequence of length *K* × *L, L* tokens are uniformly sampled to construct the input, preserving the absolute genomic coordinates of the sampled tokens. We used 50% non-contiguous data with a sampling factor *K* = 8. This strategy allows the model to capture genomic patterns across larger chromosomal regions while maintaining manageable sequence lengths.

#### Positional Encoding

We evaluate four strategies for encoding absolute chromosomal coordinates: (1) *Sinusoidal encoding* computes fixed trigonometric embeddings PE(*p*) = [sin(*p/θ*^2*i/d*^), cos(*p/θ*^2*i/d*^)] from the absolute genomic position *p*, which are added to token embeddings; (2) *ALiBi* applies linear attention biases based on scaled pairwise position differences, bias_*ij*_ = −*m* · |*p*_*i*_ − *p*_*j*_| */*10^4^, where *m* is head-specific; (3) *Linear encoding* projects the scalar coordinate to a *d*-dimensional embedding via a learnable linear layer, which is then added to token representations; and (4) *AARoPE* extends rotary position embeddings to absolute chromosomal coordinates by computing rotation matrices from genomic positions, capturing both local sequence context and long-range chromosomal interactions.

#### Analysis

As shown in Fig. 4(c), the training curves show distinct convergence patterns across positional encoding methods. While all approaches start similarly and decline rapidly in early training, AARoPE continues improving throughout the process with a consistent downward trend. In contrast, ALiBi, Linear, and Sinusoidal plateau at similar levels, indicating comparable limitations in genomic positional modeling. AARoPE also demonstrates more stable training dynamics with reduced variance, supporting its effectiveness for absolute chromosomal coordinate encoding.

### I.3 Alignment-Aware Analysis

To accommodate sequences with varying alignment availability, we design two unified input formats: **Dense** and **Dense-AA**.

#### Dense Format

For sequences without alignment information, we adopt a standardized default input format. Each token position is encoded using relative coordinates for AARoPE computation. The chromosome ID is set to 0 to indicate the absence of genomic mapping. The strand orientation is uniformly set to the forward strand (encoded as 0), and all CIGAR operations are assigned the match symbol “=” (encoded as 7).

#### Dense-AA Format

For sequences with valid alignment information, we utilize the authentic genomic coordinates and CIGAR operations obtained from sequence alignment, as described in Appendix B.5. When multiple alignments exist, we select one based on mapping quality scores during training to enhance robustness.

This dual-format design enables the model to leverage precise genomic context when available while maintaining compatibility with sequences lacking reference alignment, such as those from non-human species or novel genomic regions.

### I.4 K Sampling

USM-AA is pretrained on sequences aligned to the complete reference genome, with positional encodings derived from absolute genomic coordinates. This design enables the model to process non-contiguous input sequences, a capability that is not supported by prior models relying solely on relative positional information.

To evaluate the robustness of USM-AA under non-contiguous input settings, we introduce a *K down-sampling* strategy, controlled by a parameter *K* that quantifies the degree of sequence discretization. Specifically, given an original contiguous sequence of length *K* × *L*, we uniformly sample *L* tokens to construct an input sequence of fixed length *L*. Under this setting, the model observes only a 1*/K* fraction of the original sequence, while the absolute genomic coordinates of the sampled tokens are preserved.

For sequences containing genomic variants such as SNVs or indels, we additionally retain all variant-associated tokens together with their original genomic coordinates, and append them to the uniformly sampled tokens. Furthermore, special boundary tokens (BOS and EOS) are added to the beginning and end of each input sequence. As a result, the final input length presented to the model may exceed *L*.

#### Non-contiguous Format

To evaluate the model’s ability to handle non-contiguous input sequences, we set *K* = 8 and tested USM-AA on the NT benchmark datasets under this downsampling scheme.

As reported in Table 20, the model is capable of processing discretely sampled sequences, although its performance shows a moderate decrease compared to inputs in the Dense-AA format.

**Table 20:**
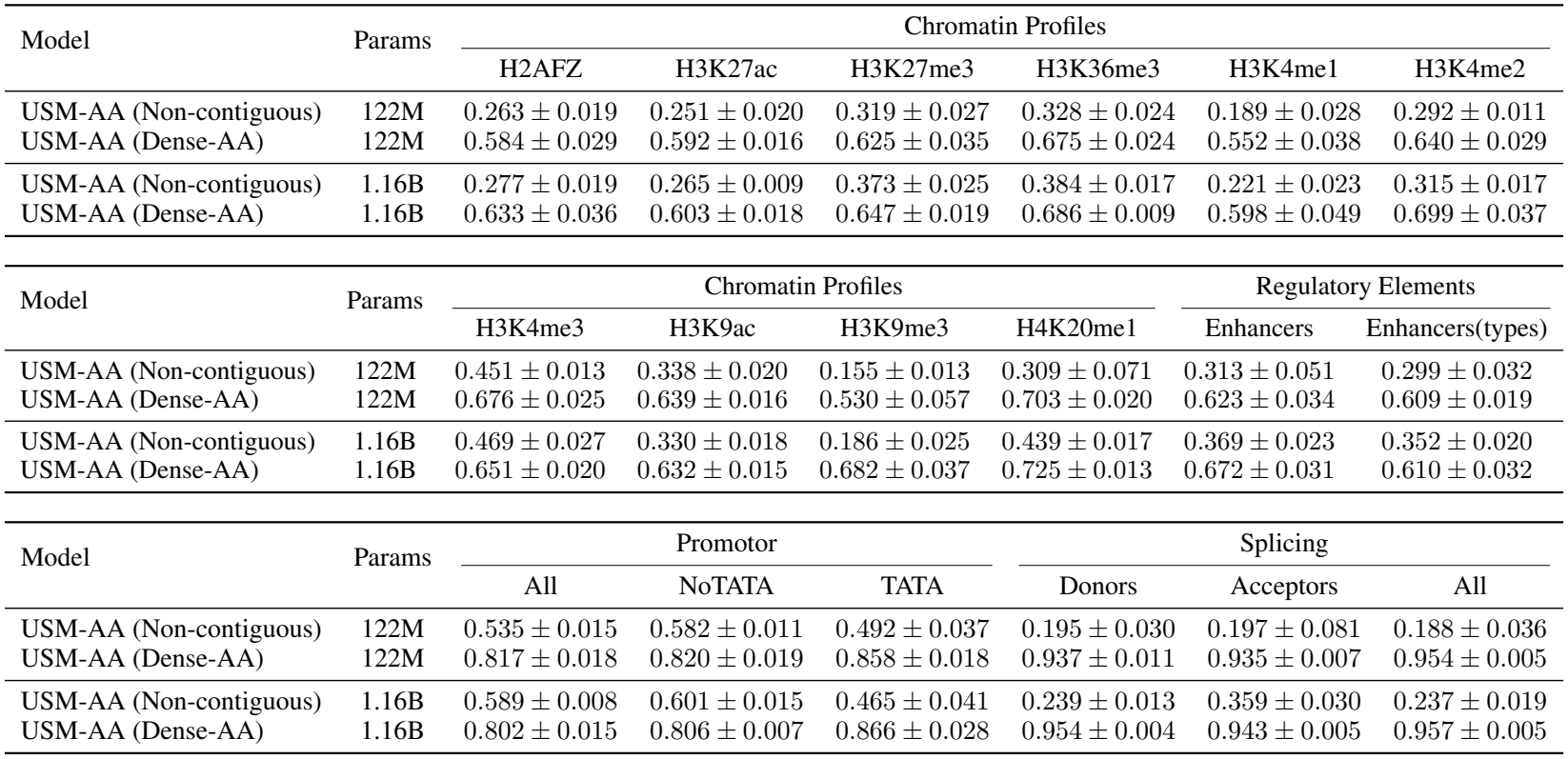
MCC performance comparison between Non-contiguous and Dense-AA input formats on NT downstream tasks.

| Model | Params | Chromatin Profiles |  |  |  |  |  |
|---|---|---|---|---|---|---|---|
|  |  | H2AFZ | H3K27ac | H3K27me3 | H3K36me3 | H3K4me1 | H3K4me2 |
| USM-AA (Non-contiguous) | 122M | 0.263 ± 0.019 | 0.251 ± 0.020 | 0.319 ± 0.027 | 0.328 ± 0.024 | 0.189 ± 0.028 | 0.292 ± 0.011 |
| USM-AA (Dense-AA) | 122M | 0.584 ± 0.029 | 0.592 ± 0.016 | 0.625 ± 0.035 | 0.675 ± 0.024 | 0.552 ± 0.038 | 0.640 ± 0.029 |
| USM-AA (Non-contiguous) | 1.16B | 0.277 ± 0.019 | 0.265 ± 0.009 | 0.373 ± 0.025 | 0.384 ± 0.017 | 0.221 ± 0.023 | 0.315 ± 0.017 |
| USM-AA (Dense-AA) | 1.16B | 0.633 ± 0.036 | 0.603 ± 0.018 | 0.647 ± 0.019 | 0.686 ± 0.009 | 0.598 ± 0.049 | 0.699 ± 0.037 |
| Model | Params | Chromatin Profiles |  |  |  | Regulatory Elements |  |
|  |  | H3K4me3 | H3K9ac | H3K9me3 | H4K20me1 | Enhancers | Enhancers(types) |
| USM-AA (Non-contiguous) | 122M | 0.451 ± 0.013 | 0.338 ± 0.020 | 0.155 ± 0.013 | 0.309 ± 0.071 | 0.313 ± 0.051 | 0.299 ± 0.032 |
| USM-AA (Dense-AA) | 122M | 0.676 ± 0.025 | 0.639 ± 0.016 | 0.530 ± 0.057 | 0.703 ± 0.020 | 0.623 ± 0.034 | 0.609 ± 0.019 |
| USM-AA (Non-contiguous) | 1.16B | 0.469 ± 0.027 | 0.330 ± 0.018 | 0.186 ± 0.025 | 0.439 ± 0.017 | 0.369 ± 0.023 | 0.352 ± 0.020 |
| USM-AA (Dense-AA) | 1.16B | 0.651 ± 0.020 | 0.632 ± 0.015 | 0.682 ± 0.037 | 0.725 ± 0.013 | 0.672 ± 0.031 | 0.610 ± 0.032 |
| Model | Params | Promotor |  |  | Splicing |  |  |
|  |  | All | NoTATA | TATA | Donors | Acceptors | All |
| USM-AA (Non-contiguous) | 122M | 0.535 ± 0.015 | 0.582 ± 0.011 | 0.492 ± 0.037 | 0.195 ± 0.030 | 0.197 ± 0.081 | 0.188 ± 0.036 |
| USM-AA (Dense-AA) | 122M | 0.817 ± 0.018 | 0.820 ± 0.019 | 0.858 ± 0.018 | 0.937 ± 0.011 | 0.935 ± 0.007 | 0.954 ± 0.005 |
| USM-AA (Non-contiguous) | 1.16B | 0.589 ± 0.008 | 0.601 ± 0.015 | 0.465 ± 0.041 | 0.239 ± 0.013 | 0.359 ± 0.030 | 0.237 ± 0.019 |
| USM-AA (Dense-AA) | 1.16B | 0.802 ± 0.015 | 0.806 ± 0.007 | 0.866 ± 0.028 | 0.954 ± 0.004 | 0.943 ± 0.005 | 0.957 ± 0.005 |

As shown in Fig. 5(a), USM-AA can still perform downstream classification on sparsely sampled non-contiguous inputs, although performance decreases as the degree of downsampling increases, reflecting the reduced sequence information available to the model.

**Figure 5:**
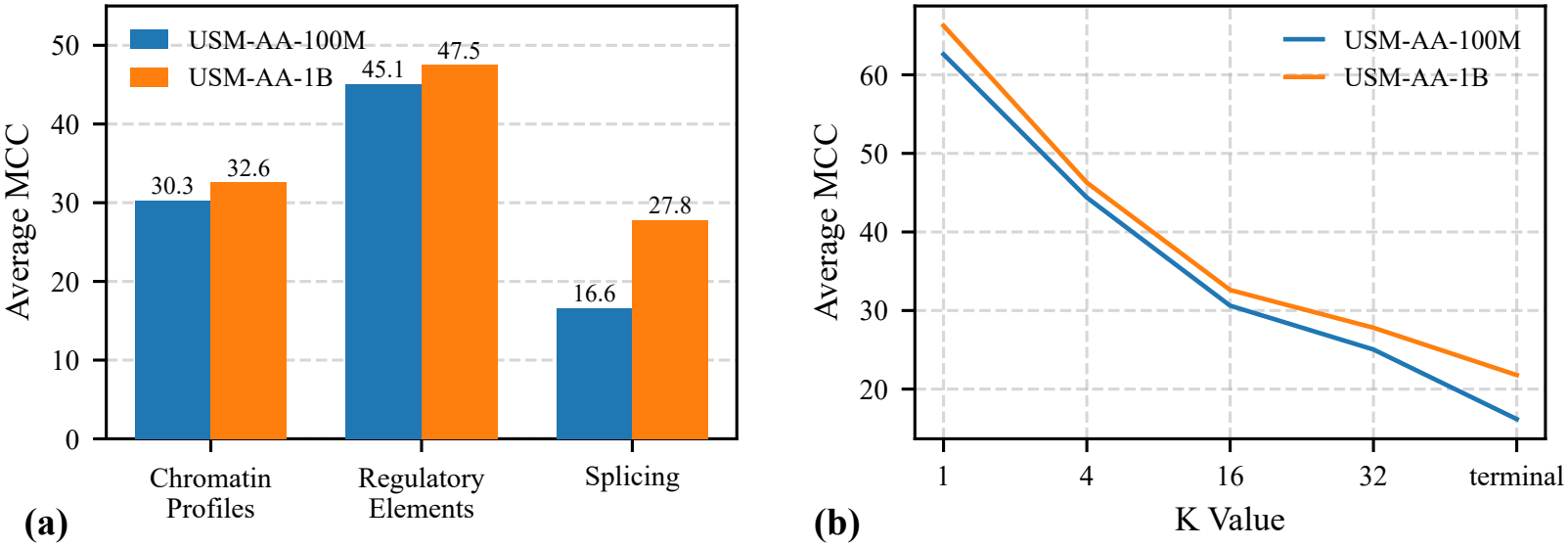
Ablation studies of USM-AA on non-contiguous input sequences. **(a)** The performance of USM-AA on non-contiguous input sequences in the NT benchmark. **(b)** Effect of different *K* values, which represent the degree of sequence sparsification, on model performance.

To further examine the effect of different sampling intervals *K* on model performance, we conduct experiments on the histone modification task from the NT benchmark, which involves relatively long input sequences. As *K* increases, the input sequences become progressively more sparse, resulting in a consistent degradation in performance, as shown in Fig. 5(b). Despite the performance decline, USM-AA maintains meaningful discriminative ability even under highly sparse input conditions.

